# Defining Quality Control Standards for Single-Cell Proteomics by Inter-Laboratory Benchmarking

**DOI:** 10.64898/2026.07.13.738155

**Authors:** Sam van Puyenbroeck, Tine Claeys, Anjali Seth, Jeewan Babu Rijal, Charline Keller, Lavender Lin, Rupert Mayer, Manuel Matzinger, Isaac Han, Pedro Aragón Fernández, Valdemaras Petrosius, Benjamin Furtwängler, Brian Boyle, Keith Rivera, Guilhem Tourniaire, Florian A. Rosenberger, Lennart Martens, Steven A. Carr, Zhen Dong, Ákos Végvári, Christine Carapito, Ryan Kelly, Karl Mechtler, Bogdan Budnik, Erwin M. Schoof, Claudia Ctortecka

**Affiliations:** CompOmics, VIB-UGent Center for Medical Biotechnology, VIB, Ghent, Belgium; Department of Biomolecular Medicine, Faculty of Medicine and Health Sciences, Ghent University Ghent, Belgium; Cellenion SASU, Lyon, France; Laboratoire de Spectrométrie de Masse BioOrganique (LSMBO), IPHC UMR7178, Université de Strasbourg, CNRS; Infrastructure Nationale de Proteomique ProFI UAR2048, Strasbourg, France; Department of Chemistry and Biochemistry, Brigham Young University, Provo, UT, USA; Research Institute of Molecular Pathology (IMP), Vienna BioCenter (VBC), Vienna, Austria; Institute of Molecular Biotechnology (IMBA), Vienna BioCenter (VBC), Vienna, Austria; Gregor Mendel Institute of Molecular Plant Biology (GMI), Vienna BioCenter (VBC), Vienna, Austria; Wyss Institute, Boston, MA, USA; Technical University of Denmark, Lyngby, Denmark; Broad Institute of MIT and Harvard, Cambridge, MA, USA; Department of Medical Biochemistry and Biophysics, Science for Life Laboratory, Karolinska Institutet, Stockholm, Sweden; Westlake University, China; Laboratory of Experimental Oncology and Radiobiology, Amsterdam UMC, Amsterdam, The Netherlands

## Abstract

Single-cell proteomics can quantify thousands of proteins from individual mammalian cells, yet the absence of community-wide quality control limits biological interpretability. Here, the HUPO Single Cell Initiative presents the first inter-laboratory single-cell proteomics benchmarking study across seven laboratories using standardized 384-well plates acquired on Orbitrap Astral and timsTOF Ultra2 instruments. Centralized analysis across six DIA software tools revealed that software choice impacts identification depth and quantitative accuracy more than instrument vendor. Multi-layered quality control enabled the detection of cell-leakage during sorting, LC misconfiguration, column degradation and site-specific pipetting failures. Inter-lab quantitative correlations were strongest between instruments of the same vendor relative to cross-platform comparisons. Sequential correction for plate identity and well position recovered clean cell-type separation for confident downstream differential expression analysis. This study provides a data-driven quality control framework spanning plate design to batch correction for reproducible single-cell proteomics across laboratories and platforms.

## Main

Multidisciplinary experimental and computational advances allow for the quantification of thousands of proteins from individual mammalian cells. Mass spectrometry-based single cell proteomics (SCP) characterizes molecular heterogeneity that imprints cell identity, state and signaling^1–7^. Unlike single-cell sequencing approaches that have identified previously unknown cellular sub-populations and transition stages^1,2^, the proteome can currently not be directly amplified, resulting in constrained sample input and dynamic range-dependent sampling biases^8,9^. This has driven rapid innovation both in sample preparation workflows to reduce potential sources of irreversible sample loss and in sensitive instrumentation over the past decade^10–20^. These methodological advances have enabled the transition from proof-of-concept demonstrations to a growing range of biological applications in which quantitative rigor is essential, including primary cells, and patient-derived samples^21–23^. Yet measurements at the picogram scale are disproportionately susceptible to contamination, carryover, batch effects, and technical noise^3–5^. Any one of these can introduce systemic biases that are difficult to distinguish from biological heterogeneity in the absence of the appropriate controls. Without common benchmarks, including controlled reference samples it is impossible to directly compare results across studies, assess the reliability of reported findings, or classify data quality.

Widespread adoption of SCP by biologists and clinicians remains limited due to the lack of standardized workflows, reporting guidelines and inter-laboratory comparisons. Recognizing this need, Gatto et al. proposed in 2023 a set of initial community recommendations for performing, benchmarking and reporting SCP experiments, advocating for the inclusion of standardized controls as routine elements of every SCP workflow^24^. Here, the HUPO Single Cell Initiative (SCI) presents the first large-scale inter-laboratory study designed to empirically validate and extend these recommendations. For this, seven independent laboratories across two continents received centrally prepared, ready-to-inject SCP plates and performed platform-specific standardized liquid chromatography mass spectrometry (LC-MS). The experimental design embeds a variety of controls for all workflow steps, enabling each quality control (QC) tier to be evaluated independently. The comprehensive dataset was evaluated across six widely used software tools, allowing the separation of software-driven effects from experimental and instrument-related variation. Collectively, this study provides a data-driven foundation for experimental design suggestions and data QC recommendations for the SCP community to assess/ reproducibility and finalize clinical translation. For this, we provide (i) a raw and processed benchmarking datasets with full metadata annotation; (ii) initial software benchmarking applying developer-recommended search settings; (iii) instrument vendor-specific data evaluation; (iv) inter-/intra-laboratory variance estimations and; (v) a multi-tier SCP-specific QC framework.

## Results

### An inter-laboratory framework for standardized SCP quality control

As part of the HUPO SCI, we conducted an inter-laboratory study across seven independent laboratories designed to define and benchmark the quality control measures necessary for rigorous SCP. Decoupling sample preparation from instrument acquisition across sites allows for a direct assessment of cross-site and cross-platform reproducibility under tightly controlled pre-analytical conditions.

Central to the study is the 384-well plate layout that integrates multiple orthogonal quality control tiers alongside the single-cell samples (**Fig. 1a**). HEK293T and K562 cells were selected for their well-characterized proteomes and comparable size distributions. 108 single cells per cell type (216 total) were deposited using image-based cell isolation within the cellenONE picolitre dispensing platform. The cellenONE not only allows for all downstream sample preparation and incubation, but also includes visual inspection of all isolated cells for QC purposes. The cells were dispensed into the central wells of each plate with predeposited MasterMix including an MS compatible detergent and a dual-protease mix, as previously described^19^. Proteins were digested for 2 hours while being continuously rehydrated to minimize evaporation. The digestion was then quenched and plates with digested single-cell samples alongside seven QC tiers to isolate and monitor distinct sources of variability were sealed and frozen prior to shipment to all participating sites (**Supplemental Table 1**).

**Figure 1:**
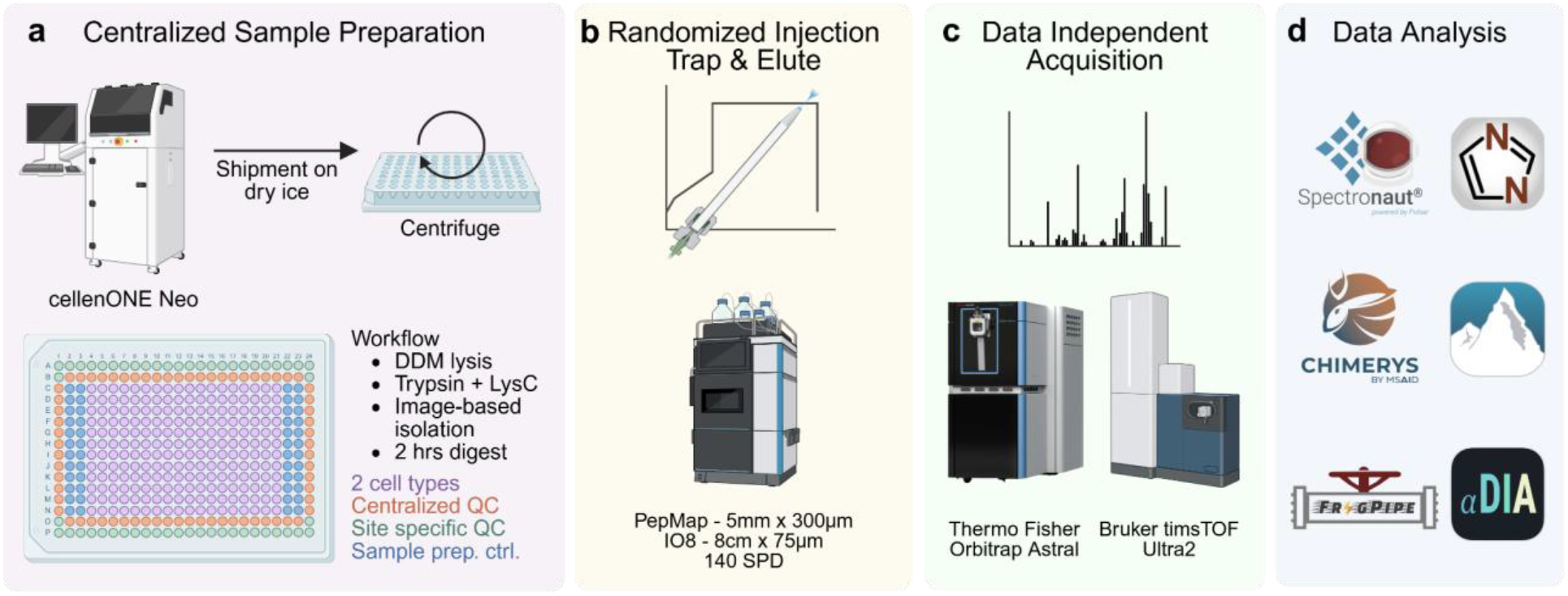
Inter-laboratory workflow outline from centralized sample generation and data analysis to standardized LC MS/MS. **(a)** The centralized sample preparation was performed using the cellenONE cell isolation and picolitre-dispensing system. 384-well plate design include centralized QC samples (a quantitative three proteome-mix in 2 different ratios and a qualitative HeLa dilution at 200 pg; n = 22 each per plate; and blanks n = 24 per plate), site-specific QCs (freshly prepared HeLa dilution and blanks; n = 24 and 28 per plate, respectively), sample preparation controls (processing control containing all buffers used during sample preparation, sorting control also contains the sorting liquid without a cell; n = 24 each per plate) and 2 different cell types (HEK293T and K562; n = 108 each per plate) that were lysed and digested in DDM with a Trypsin/Lys-C combination for 2 h at 45°C with continuous rehydration. The sealed plates were shipped to all participating laboratories, where they were centrifuged after thawing and prior to acquisition. **(b)** All samples were injected according to a uniform but randomized injection scheme, alternating QC and single-cell sample blocks on a Thermo Fisher Vanquish Neo with a PepMap 5-mm trap and an Ionopticks 8-cm analytical column. **(c)** Samples were acquired on either a Thermo Fisher Orbitrap Astral (n = 6) or a Bruker timsTOF Ultra2 (n = 3). Of note, except for one Astral using ABIRD, all other instruments were equipped with the FAIMS Pro Duo interface. **(d)** Centralized data analysis was performed using six software tools on the quantitative QC mixes for selection of optimal downstream tool, including Spectronaut, DIA-NN, CHIMERYS, PEAKS, FragPipe and alphaDIA^25–28^.

QC samples in each well plate included the following. First, the quantitative QC mix (n=22) comprised two distinct ratios of a three-proteome mixture of human, yeast and *Escherichia coli* (*E. coli*) detailed in the method section and was injected at 200 pg human material and a total of 307.7 pg per injection. This mixture allowed for an instrument-level quantitative accuracy benchmark and assessment of technical variability independent of the biological sample heterogeneity. Second, for qualitative QC (n=22) we diluted a commercially available HeLa Protein Digest Standard (200 pg per injection), providing a sensitivity reference across sites and platforms. Third, the processing control (n=24 per plate) contains equal volumes of MasterMix, rehydration and quenching solvent without a cell, while fourth, the sorting control (n=24) received a droplet of the sorting media without a cell. These dual sample processing controls allow for independent quantification of reagent and cell dispensing backgrounds. Fifth, blank wells (n=24) contained only 0.1% formic acid to monitor carryover between injections. Sixth, prior to acquisition each site followed a standardized thawing protocol and appended site-specific qualitative controls (diluted HeLa Digest to 200 pg per injection; n=24) and seventh, additional site-specific blank injections of 0.1% formic acid (n=28) to correct for any signal variability attributable to shipment or freeze-thaw cycles. A pre-randomized acquisition sequence, alternating blocks of single-cells and QC samples, was enforced across all participating sites.

All seven laboratories used an identical chromatographic setup consisting of a Vanquish Neo UHPLC system with a 5-mm PepMap trapping column in combination with an IonOpticks 8-cm analytical column, operating with a dedicated gradient to allow a throughput of 140 samples per day (SPD; **Fig. 1b**). Data were acquired across two MS platforms, including six plates on the Orbitrap Astral equipped with FAIMS Pro Duo, one Orbitrap Astral equipped with an active background ion reduction device (ABIRD; Astral_4), and three on timsTOF Ultra2 instruments (**Fig. 1c**). To disentangle instrument-level from software-level contributions to data quality, all raw data was processed centrally using six DIA analysis tools: Spectronaut (20.3), DIA-NN (2.6.0), CHIMERYS (5.0.0), PEAKS Online (13), FragPipe (24.0), and alphaDIA (2.0.1) (**Fig. 1d**)^25–28^. This multi-software strategy enables systematic evaluation of how differences in raw data processing affect reported proteome depth, quantitative accuracy, and data completeness.

### Software choice drives quantitative accuracy independent of instrument performance

The low protein yield of mammalian single cells challenges confident identification and accurate quantification. To benchmark this, we subjected all Astral and timsTOF Ultra2 datasets of the three-proteome quantitative QC mix to analysis with the six DIA software tools (**Fig. 2a**). Each tool was executed according to its developer recommended search strategy (details in Materials and Methods). The three-proteome quantitative QC mix described above, was injected at 307.7 pg to provide a ground-truth framework for evaluating whether measured fold-changes between species match their expected values.

**Figure 2:**
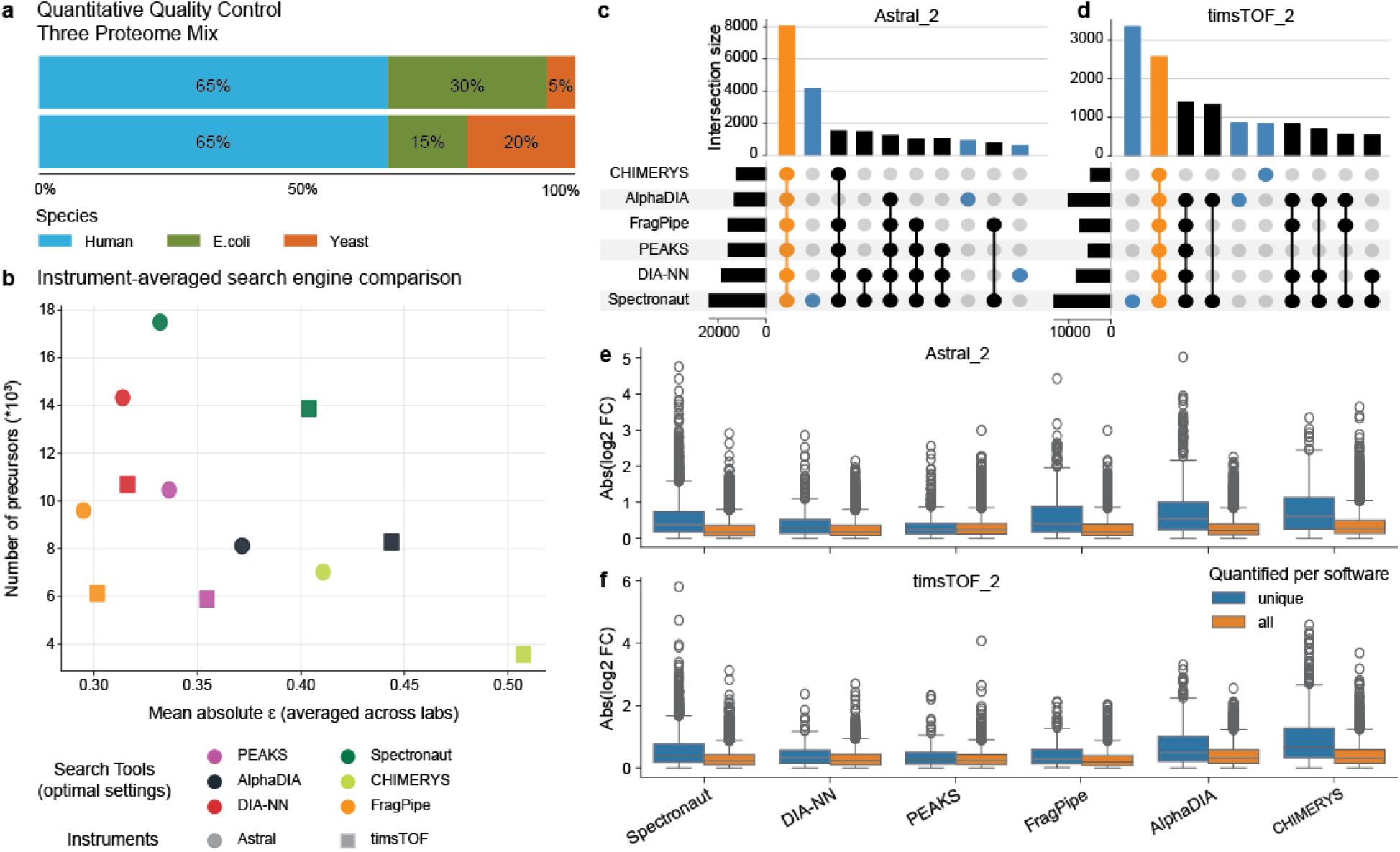
Quantitative accuracy and software comparison using a three-proteome mix at ∼300 pg input. **(a)** Relative ratios of *E. coli*, human and yeast peptides at two abundance ratios injected at 307.7 pg total input (n = 22 per laboratory site). **(b)** The average of quantified precursors detected in at least three raw files within each condition plotted against the mean absolute difference between measured and expected log_2_ fold change averaged for six Astral and three timsTOF Ultra2 instruments. Each point represents the average of the instrument–type to software combination; color indicates search engine and symbol indicates instrument vendor. **(c-d)** Peptide sequence overlap across all six search tools for the representative (c) Astral_2 and (d) timsTOF Ultra2_2 instrument; blue bars indicate tool-unique identifications, orange indicates sequences identified by all six tools. **(e-f)** Absolute log_2_ fold-change error for peptides identified exclusively by a single tool (blue) versus peptides shared across all tools (orange) for (e) Astral_2 and (f) timsTOF Ultra2_2.

To directly connect depth with accuracy, we combined the number of quantified precursors detected in at least three raw files in each condition with the mean absolute difference of measured and expected fold-changes. Across all instruments, we found that software choice is a primary determinant of both identification depth and quantitative accuracy, exceeding the contribution of instrument identity within each platform (**Fig. 2b; Supplementary Fig. 1a-b**). Across the six Astral instruments and all search engines, precursor identification count ranged from 4,642 to 24,889, while the timsTOF Ultra2 instruments the range spanned 3,104 to 14,686. Spectronaut consistently yielded the highest number of identified precursors on both platforms with higher quantitative errors compared to the second best performing tool, DIA-NN. FragPipe and PEAKS showed intermediate identification numbers for both instruments, while alphaDIA is the only tool recovering similar unique precursors in Astral and timsTOF Ultra2 instruments (**Fig. 2b**). While FragPipe has the lowest quantitative error of all the tools, identification depth is reduced (**Fig. 2b; Supplementary Fig. 1a-b**). Overall, on the timsTOF Ultra2, CHIMERYS yielded the least accurate quantification combined with relatively shallow identifications.

During initial comparative analysis, a systematic offset in log_2_ space was observed for PEAKS Online and CHIMERYS that was not attributable to sample or instrument effects. This deviation was corrected by applying run-specific total ion current (TIC) normalization prior to fold-change calculation, which resolved the bias compared to all other tools (**Fig. 2d; Supplementary Fig. 1b**). Due to the application of DIA-NN quantification in FragPipe workflows, their per-species fold-change distributions normalized by precursor count consistently reveals a nearly perfect overlap across both software tools. Spectronaut and DIA-NN show comparable distribution widths and centering for the Astral dataset, with Spectronaut showing a slight broadening on the timsTOF Ultra2. This, we attribute in part to the substantially greater number of precursors Spectronaut identifies in timsTOF datasets relative to the other tools and is independent of their relative abundance (**Extended Fig. 1; Extended Fig. 2**). Interestingly, the quantitative error for the Astral versus timsTOF instruments was comparable for DIA-NN, FragPipe and PEAKS, with larger variability for Spectronaut, CHYMERIS and alphaDIA (**Fig. 2b**). These differences likely reflect a combination of vendor-specific signal processing and tool-dependent quantification algorithms^6,7^. While the implementation of FAIMS Pro Duo aims at reducing singly charged precursor ions to reduce spectral complexity, Astral_3 and Astral_5 exhibited the highest inter-tool variability. This is despite Astral_4 being the only instrument in the cohort operated without FAIMS Pro Duo (**Fig. 2d**). This indicates the quantitative accuracy is primarily influenced by abundance, but also software and instrument dependent as opposed to the suspected instrument vendor impact.

To assess the composition and confidence of tool-specific identifications, we examined the overlap of identified peptide sequences across all six software tools within the representative Astral_2 and timsTOF Ultra2_2 (**Fig. 2e-f; Extended Fig. 3a-g**). Of note, the samples acquired on Astral_6 underwent extended pre-analytical storage and its *quantitative QCs* were thus not included in the search tool comparison. The intersection shared across all six tools, highlighted in orange, represents the highest-confidence core peptides, comprising 7,530 sequences for the representative Astral and 2,375 for the timsTOF Ultra2. Spectronaut contributed the largest number of tool-unique identifications on both platforms, while showing the greatest overlap with alternative tools. This suggests that Spectronaut’s additional identifications are not isolated marginal detections but could represent extensions of otherwise unidentified peptides. To confirm this, we compared the quantitative error of peptides identified exclusively by a single tool against those identified by all tools (**Fig. 2f-g; Extended Fig. 4a-g**). As expected, peptides shared across all software consistently showed the smallest quantitative error, while peptides uniquely identified by CHIMERYS showed the largest median error across both instrument platforms. Despite FragPipe’s use of DIA-NN for quantification, peptides uniquely identified by FragPipe exhibited a larger error margin than those uniquely identified by DIA-NN. This suggests that the additional identifications made by FragPipe’s pseudo DDA search strategy are enriched for lower-confidence detections^8,9^. Spectronaut’s large pool of unique identifications showed a comparable median error relative to other tools, but with the greatest number of outliers for both instruments (**Fig. 2f-g; Extended Fig. 4a-g**). Interestingly, examination of the physicochemical properties of tool-specific unique peptides revealed broadly comparable characteristics, including peptide length, precursor charge, missed cleavages and even sample-preparation specific post-translational modifications (PTMs), like oxidation (**Supplementary Fig. 2a-h**). Together this data suggests that Spectronaut directDIA analysis identifies the highest number of high confidence peptides and was thus selected as the primary processing tool for all downstream single-cell analyses reported in this study. Importantly, this selection does not preclude the use of alternative tools; rather, based on the quantitative QC samples, it reflects the optimal combination of identification depth and quantitative accuracy within the present dataset. For transparency and reproducibility, all downstream analyses were repeated using DIA-NN, with all figures made publicly available on GitHub. Importantly, all trends reported here for Spectronaut were recapitulated in the DIA-NN dataset, confirming that the key findings of this study are not software-specific.

### Identifying and quantifying the contribution of variability from sample preparation to data acquisition

To estimate the total variability observed in SCP data we aim to deconvolve the contributions of instrument performance, site-specific handling, and sample preparation background. For ease of interpretation, we solely present Spectronaut directDIA data for comparisons of technical variability across instruments, sample types, and QC tiers. Importantly, relative to Spectronaut DIA-NN resulted in more IDs for single-cell samples on Astral instruments and lower IDs on timsTOF data, while the quantitative accuracy and all additional data trends presented in the manuscript are conserved across tools (**Supplementary Fig. 7**; DIA-NN data available on GitHub). Additionally we want to highlight the power of the free DIA-NN utilization for academic groups for Astral instruments, while Spectronaut analysis shows a significant advantage for timsTOF data.

Across all nine instruments, quantified precursors and protein groups varied substantially even within sample types (**Fig. 3a-b**). Astral_2 yielded the highest median precursor and protein identifications from *single cells* (10,706 precursors; 2,850 protein groups) but also exhibited the largest within-instrument spread across both instrument platforms. Interestingly, this pattern was mirrored in its 200 pg QC samples, which in combination with the high intensities detected in the control samples, suggests that the additional precursor identifications in part reflect increased sampling of sorting background signal rather than a uniform improvement in sensitivity (**Fig. 3d-e; Supplementary Fig. 8**). This is while most *blank injections*, both centralized and site-specific, yielded near-zero identifications across instruments (**Fig. 3a-b**). The *centralized blanks* confirm the sealed-plate shipment integrity and the site-specific blanks indicate the absence of carryover. Based on this, the comparable identifications between HEK293T and K562 cells across all instruments confirm that the choice of two similarly sized cell types did not introduce a systematic size-driven asymmetry into the datasets.

**Figure 3:**
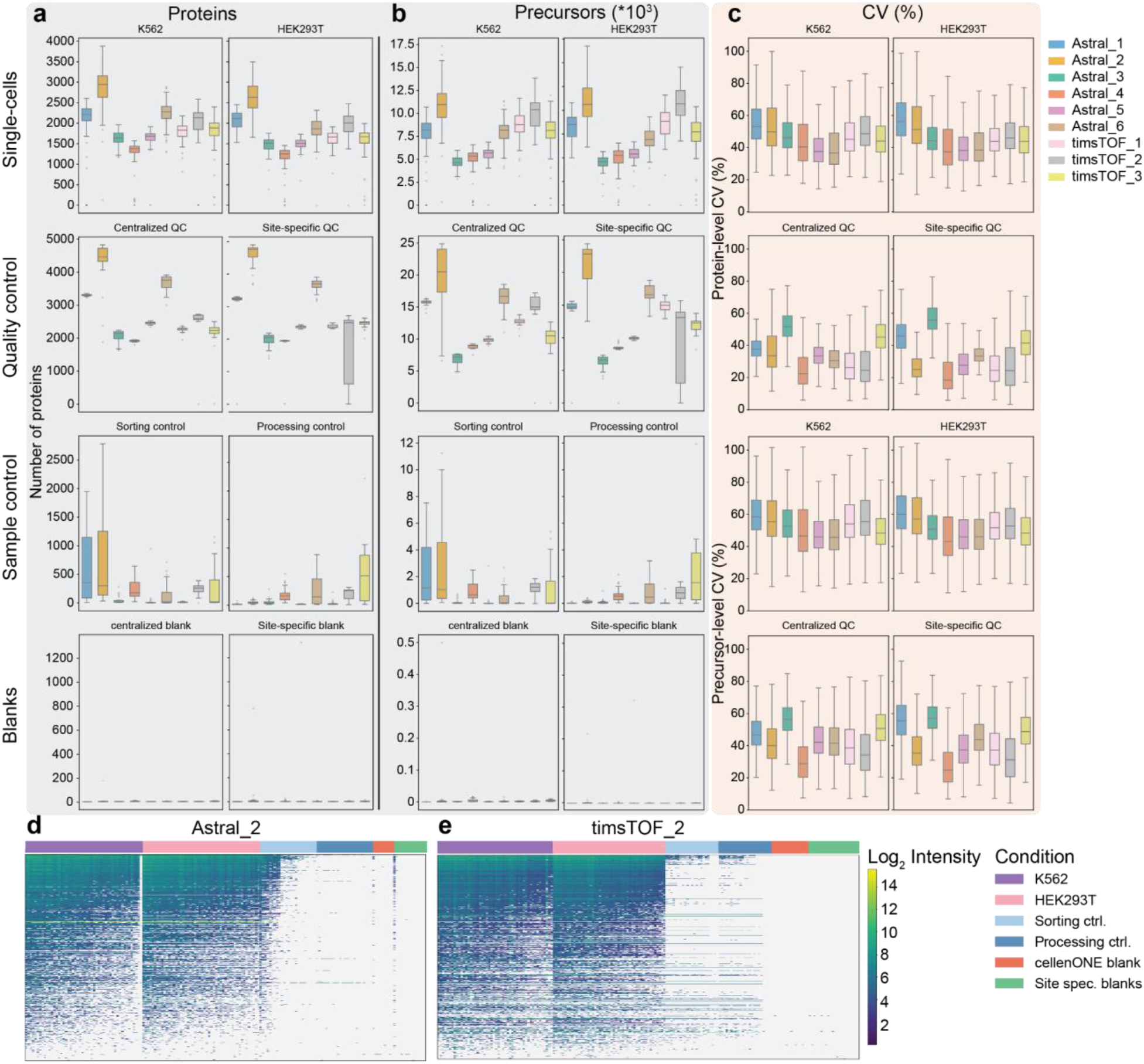
Quantitative and qualitative performance across all instruments and sample types. Number of quantified **(a)** precursors and **(b)** proteins per sample across all instruments, shown for HEK293T and K562 single cells, centralized HeLa QC, site-specific HeLa QC, processing controls, sorting controls, cellenONE blanks, and site-specific blanks. **(c)** Protein-level (top) and precursor-level (bottom) %CV for the same sample categories as a-b displayed as violin plots. Overlap of protein identifications between single-cell samples (HEK293T, purple; K562, pink), processing controls (light blue), sorting controls (dark blue), and blank injections (orange and green) for a representative **(d)** Astral_2 and **(e)** timsTOF Ultra2_2 plate. Heatmaps for all other instruments are displayed in Supplementary Fig. 7. Blue cells indicate identified proteins; grey cells indicate missing values.

The *centralized QC* sample showed tight identification distributions across most instruments, with a median coefficient of variation (CV) ranging from approximately 23.4 % (Astral_4) to 52.3 % (Astral_3) at the precursor level (**Fig. 3c**). Astral_4, notably the only Astral instrument in this study operated without FAIMS and instead using ABIRD, produced the tightest QC identification distributions and the lowest quantitative CVs across both centralized and site-specific HeLa samples (precursor median CV: 23.4 % centralized, 19.1 % site-specific; protein median CV: 21.3 % centralized, 18.6 % site-specific; **Fig. 3c**). This indicates that, for clean peptide standards at 200 pg input amounts, IM-based background suppression is not a prerequisite for high quantitative reproducibility as other sources of chemical noise are absent. Among the timsTOF instruments, timsTOF_2 showed the largest discrepancy between centralized and *site-specific QC* CVs, which we attribute to variability introduced during local sample preparation at that site, consistent with the rationale for including site-specific QC samples as a shipment and handling monitor.

The *processing and sorting controls* revealed instrument- and plate-specific differences in sample preparation background that would be invisible without their inclusion (**Fig. 3a-b**). Astral_2 showed minimal signal in its processing control but the highest signal among all instruments in the sorting control, indicating that the cell suspension used for this plate may have contained a proportion of lysed or leaking cells prior to dispensing. Because all identifications in these controls were filtered against a standard contaminant database (i.e., including common environmental contaminants, cell culture media components, and keratins) the peptides detected represent genuine intracellular protein material introduced as background that are also present in the single-cell samples (**Fig. 3d-e; Extended Fig. 6a-g**). The elevated single-cell identifications in Astral_2 may therefore reflect systematic inflation from cell-leakage background rather than instrument sensitivity alone (**Fig. 3d**). In contrast, Astral_4, timsTOF_2, and timsTOF_3 showed signal in both processing and sorting controls but clean blanks, localizing the source of background to the sample preparation steps rather than instrument-level carryover or contamination (**Fig. 3e; Extended Fig. 6c, g**). Proteins detected in control samples cannot be reliably attributed to single-cell biology without correction for background levels (**Fig. 3d-e; Extended Fig. 6a-g**). We therefore recommend that downstream analyses explicitly cross-reference identified proteins against the corresponding processing and sorting control identifications.

Focusing on the *single-cell samples* specifically, precursor-level median CVs across all instruments ranged from approximately 38.6 % to 57.5 % for both HEK293T and K562 cells (**Fig. 3c**). The two cell types showed closely matched CV distributions within each instrument, reinforcing the consistency of the centralized preparation. Astral_5 produced the tightest single-cell CVs (39.9 % HEK293T, 39.6 % K562 at precursor level), while Astral_1 showed the highest (57.5 % and 54.3 % at precursor level). All three timsTOF Ultra2 instruments yielded precursor identifications broadly comparable to Astral_1, Astral_2, and Astral_6, but matched the protein-group identification depth of all Astral instruments except Astral_2, which yielded approximately 54.4 % more protein groups than the indicated instruments (**Fig. 3a-b**). Importantly, even the 200 pg centralized QC samples showed precursor-level CVs in the range of 23-52 % across instruments (**Fig. 3c**). The single-cell input level specific multi-tier QC design provides orthogonal references at each level of the workflow to systematically flag outliers and attribute variability to sample preparation, instrument performance, or biological differences between cell types.

### Establishing quality thresholds and detecting acquisition failures across all layers of the workflow

Beyond global identification and CV metrics, robust SCP workflows require practical quality control criteria that can identify failed injections, instrument decline, and sample preparation anomalies in real-time or during post-acquisition review. For this we systematically evaluated *chromatographic stability* across the gradient, spatial distribution of identifications across the plate, identification consistency over the course of the acquisition, and overall data completeness. As a first-pass filter, runs retaining fewer than 10% of the median identification count for their sample type should be flagged for additional review or exclusion. For this we evaluated the instrument-specific differences in the proportion of precursor identifications per retention time (RT) bin (**Fig. 4a-b**). The QC and single-cell elution profiles serve as sensitive indicators of chromatographic performance and reproducibility (**Fig. 4a-b; Extended Fig. 7a-i**). Importantly, Astral_4 contained a subset of injections with dramatically aberrant elution profiles that deviated markedly from the expected pattern (**Extended Fig. 7d**). Cross-referencing with the metadata confirmed that these corresponded to the first injection block of that plate, during which an incorrect percentage of solvent B was applied. These injections were subsequently excluded from all downstream analyses, demonstrating that RT stability is an effective tool for the detection of chromatographic differences that could influence relative quantification.

**Figure 4.**
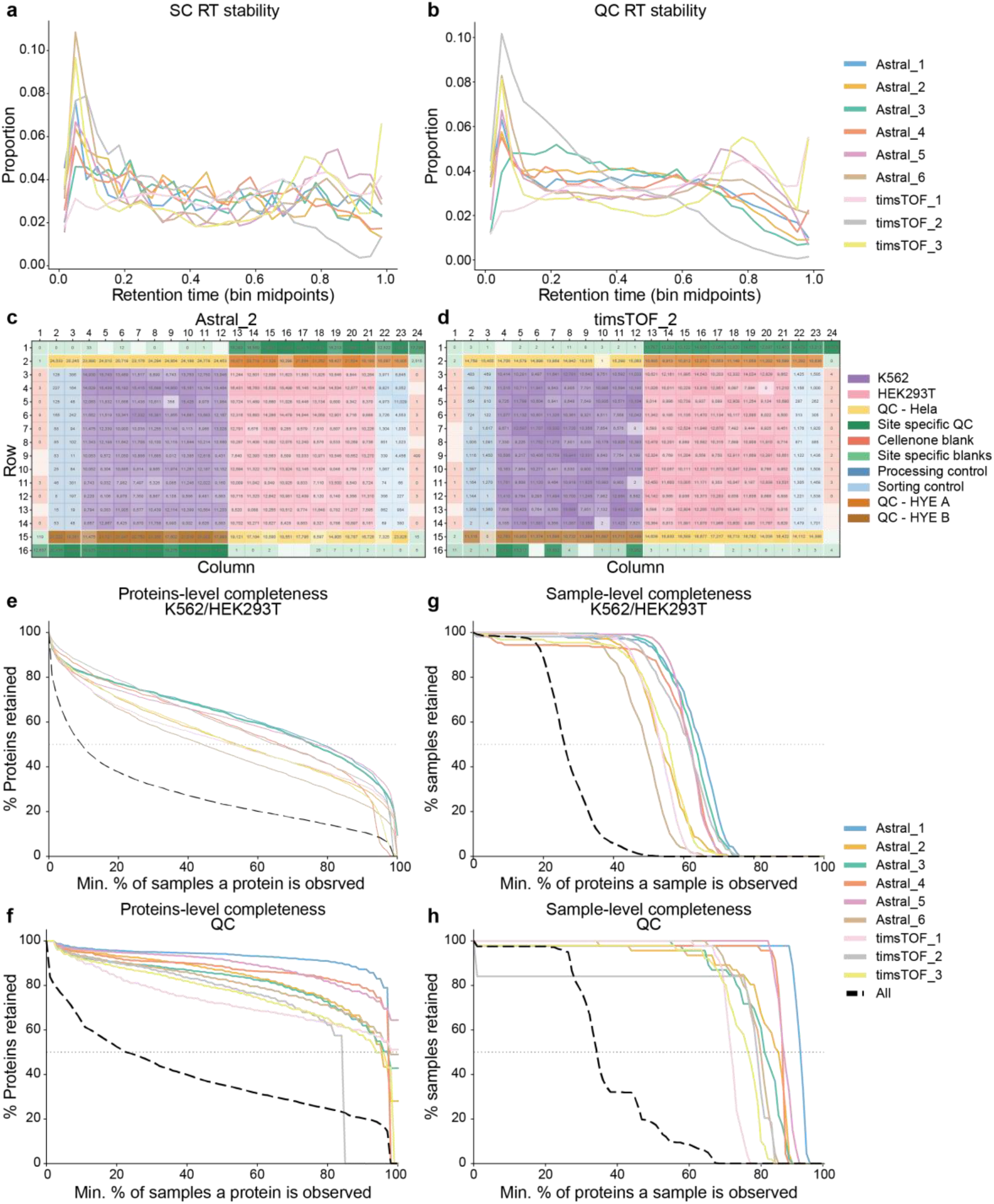
Multi-tier quality control evaluation of single-cell acquisitions. Percentage of precursor identifications across normalized retention time (RT) bins for **(a)** single-cell samples and **(b)** QC samples across all nine instruments (Astral, n = 6; timsTOF Ultra2, n = 3). 384-well plate overview showing precursor identifications per well for representative **(c)** Astral_2 and **(d)** timsTOF Ultra2_2 plates; colors indicate sample group and color intensity reflects relative identification counts within each group. Feature-level data completeness showing the minimum percentage of samples in which a protein must be observed versus the percentage of all proteins retained, for **(e)** single-cell samples and **(f)** HeLa QC samples. Sample-level data completeness showing the minimum percentage of proteins a sample must contain versus the percentage of samples retained, for **(g)** single-cell samples and **(h)** HeLa QC samples.

Next, we aimed to evaluate regular *edge effects* across the 384-well plate through visualizing precursor identifications as a spatial heatmap (**Fig. 4c-d; Extended Fig. 8a-g**). Rather than systematic preparation or evaporation effects, identification dropouts were distributed as isolated single-well events spread stochastically across the plate. As expected, the low-identification wells in Astral_4 co-localized precisely with the injection positions identified by the RT stability analysis as affected by incorrect solvent B composition (**Extended Fig. 7d; Extended Fig. 8c**). For timsTOF_2, the elevated variability in site-specific QC samples was spatially localized to one region of the plate, consistent with a pipetting or irregularity during local QC sample addition rather than instrument performance. To confirm that longitudinal instrument performance was stable across instruments, we correlated injection order with identified precursors (**Extended Fig. 9a-i**). Globally, the QC samples tracked the same trends as the single cells within each instrument, confirming that instrument-specific decline can be separated from sample dropouts. Astral_1, Astral_2, Astral_5, timsTOF_1, and timsTOF_2 all showed a gradual decline in the later acquisitions relative to the earlier ones, consistent with minor column aging or progressive accumulation of chemical background under high-throughput conditions (**Extended Fig. 9a, b, e, g, h**). Astral_3 displayed an initial increase followed by a subsequent decline, potentially reflecting column conditioning in the early acquisition phase (**Extended Fig. 9c**). In contrast, Astral_4 showed an increase in identification numbers from early to late acquisitions across all sample types, which in context represents recovery from the low-identification first block caused by an unconditioned analytical column (**Extended Fig. 9d**).

The fraction of precursors or proteins quantified across samples is a critical determinant of downstream statistical power and biological interpretability. As expected, *feature-level completeness* curves showed a sharp initial drop-off for single cells across all instruments, with many precursors detected in only one or two samples (**Fig. 4e-f**). Astral_1, Astral_3, and Astral_5 retained the highest proportion of precursors and proteins across increasing completeness thresholds, followed by Astral_4 and timsTOF_2. Astral_2 and Astral_6, despite yielding high absolute identification numbers in individual samples, showed relatively poor feature-level completeness. At the sample level, most instruments retained close to 100% of single-cell samples when requiring detection in fewer than approximately 40–50% of precursors, with a progressive decline thereafter (**Fig. 4g-h**). QC samples showed substantially steeper and higher completeness curves than single cells on all instruments, confirming more reproducible sampling of clean 200 pg input (**Extended Fig. 10a-b**). Interestingly, timsTOF_2 indicated an immediate drop in sample-level completeness only within QC samples, complementary confirming the pipetting or injection failures located in column 16 within the plate (**Fig. 4d, g, h; Extended Fig. 10a-b**).

Our suggested four layers of QC allow for systematic identification of chromatographic failure and distinction of failed sample preparation (i.e., cell isolation as also annotated in the metadata or identifications within processing and sorting controls) from possible plate-location- or injection-count-dependent biases.

### Intra- and inter-laboratory quantitative reproducibility is instrument-specific and vendor-dependent

Intra-instrument quantitative reproducibility independently of cellular variability allows estimation of the expected technical and biologically relevant variation. For this, we computed pairwise Pearson correlations between single cells of the same type within each instrument (**Fig. 5a-b**). Sorting instruments by median *intra-lab correlation*, timsTOF_2 led for both HEK293T (∼0.93) and K562 (∼0.92), followed by Astral_1 and Astral_2. The instruments with the lowest intra-lab correlation for single cells, Astral_3 Astral_6, and timsTOF_3, still maintained median Pearson correlations of approximately 0.90. This demonstrates that even the lower-performing instruments in this cohort achieve excellent quantitative SCP reproducibility at 140 SPD and one-pot sample preparation. Notably, Astral_3 and Astral_6 are physically distinct instruments housed in the same laboratory, indicating that the observed variability within single-cells and QC samples is instrument-specific rather than site-specific (**Fig. 5a-d**). Additionally, the ranking shifted when examining QC samples, where Astral_1 led for site-specific QC (∼0.962 median), while timsTOF_2 dropped relative to its single-cell ranking, consistent with the site-specific preparation variability identified earlier. Astral_2 showed a marked decline in correlation for the centralized HeLa QC relative to its single-cell performance. This confirms comparable quantitative performance of QC 200 pg peptide input or cell-line samples and highlights the importance of in-depth evaluation of the multi-tier QCs.

**Figure 5:**
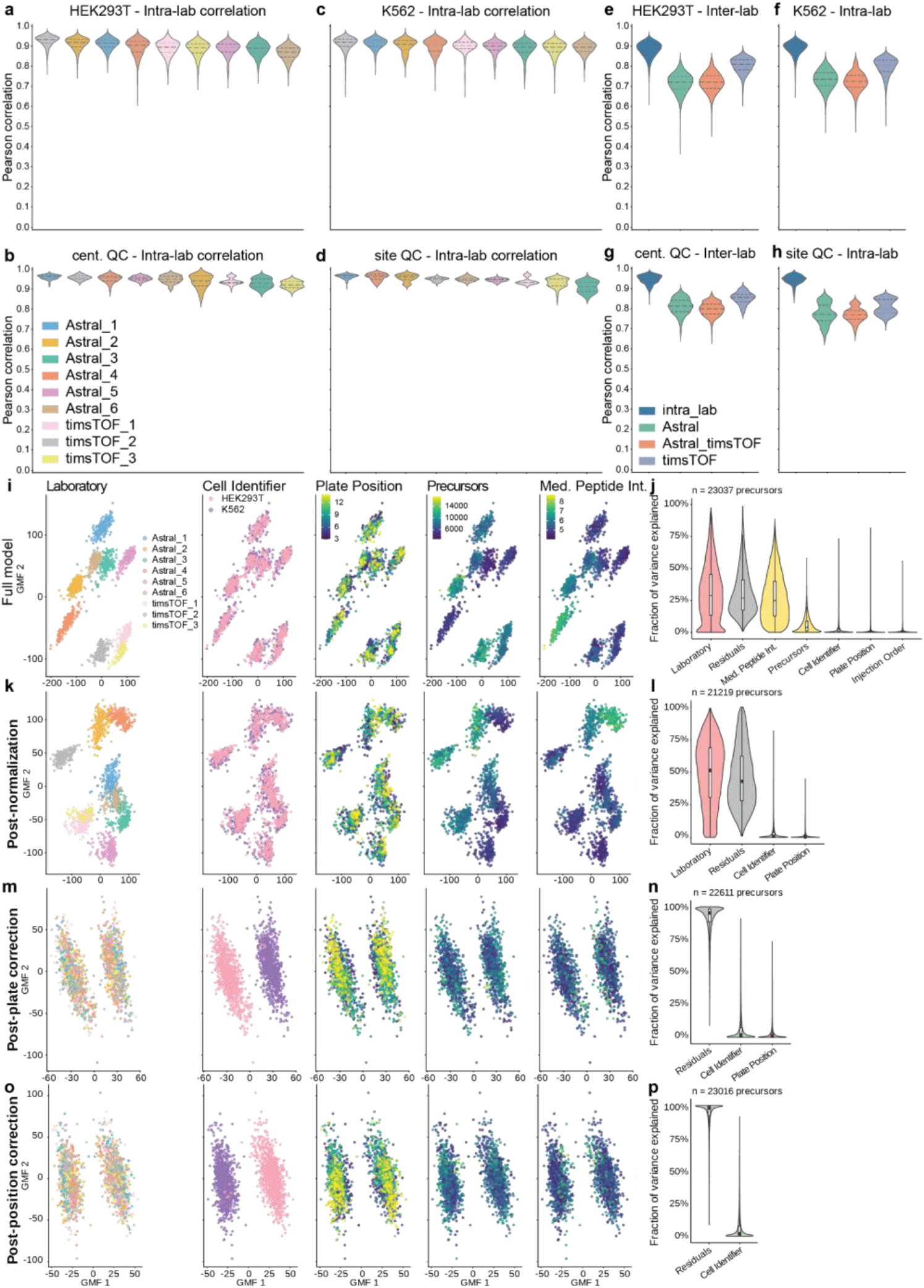
Intra- and inter-laboratory quantitative reproducibility across single-cell and QC samples. Violin plots of pairwise intra-laboratory Pearson correlations ranked by median for **(a)** HEK293T single cells, **(b)** K562 single cells, **(c)** site-specific HeLa QC, and **(d)** centralized HeLa QC across all nine instruments (Astral, n = 6; timsTOF Ultra2, n = 3). Inter-laboratory Pearson correlations stratified by instrument comparison type, intra-lab (dark blue), Astral vs. Astral (green), Astral vs. timsTOF Ultra2 (orange), timsTOF Ultra2 vs. timsTOF Ultra2 (purple) for **(e)** HEK293T single cells, **(f)** K562 single cells, **(g)** site-specific HeLa QC, and **(h)** centralized HeLa QC. omicsGMF latent factor representation colored by (from left to right) participating laboratory, cell identity, location within the plate, number of precursors and median intensity for **(i)** uncorrected, (k) normalized **(m)** plate ID or **(o)** plate ID and plate localization corrected precursor quantities with the respective **(j-n)** fraction of variance explained.

Expanding the correlation analysis to all pairwise instrument combinations revealed a clear hierarchy of reproducibility similar for single-cell samples and 200 pg peptide input (**Fig. 5e-h**). *Intra-lab correlations* were the highest across all sample types (median ∼0.90 for single cells), reflecting the similarity of technical/background noise and instrument-specific variation when comparing samples prepared and acquired under identical local conditions. Cross-instrument correlations dropped substantially for comparisons between Astral instruments (0.73 for single-cells and 0.78 for QC samples), between Astral and timsTOF instruments (0.73 for single-cells and 0.77 for QC samples), or between timsTOF instruments (0.82 for single-cells and 0.83 for QC samples) (**Fig. 5e-h**). We hypothesized that the higher median correlation of timsTOF-to-timsTOF comparisons was likely influenced by the smaller number of timsTOF instruments in this cohort. However, a pairwise correlation of individual instruments revealed marginally better agreement between timsTOF instruments (0.77-0.84 across single-cell and QC samples), compared to Astral instruments operated with FAIMS (0.68-0.85) entirely separating Astral_4 in the analysis of single-cells (0.65-0.75**; Supplementary Fig. 3a-d**). This confirms that distinct background removal strategies drive quantitative divergence more drastically in complex single-cell compared to clean QC samples. To the best of our knowledge, this study is the first to evaluate cross-instrument-vendor variability on the same sample, demonstrating that instrument vendor harmonization alone is insufficient to achieve high inter-laboratory reproducibility without additional normalization strategies.

Next, we determined how much of the observed variance in single-cell precursor intensities can be assigned to technical sources, like plate identity, acquisition site, and well position, versus biological signal. For this we applied variance partitioning analysis at the precursor level using scplainer, to dissect total variance into respective contributions from defined technical and biological covariates^10^. This was complemented with decomposition using generalized matrix factorization through omicsGMF^11^ of precursor intensities revealed plate identity as the primary source of variance, which persisted after normalization (**Fig 5i-k**). This approach handles missing values natively, allowing a low-dimensional latent representation to be built at each correction stage without prior imputation or transformation. Within the omicsGMF latent factor representation, instrument platform type was a core separating feature, with Astral and timsTOF Ultra2 instruments forming distinct clusters. Within the Astral group, Astral_2 (highest overall identifications) and Astral_4 (Astral-ABIRD) are separated further from the others. In the uncorrected and normalized latent factor representation, the two cell types showed complete overlap which confirms that technical variance from plate and instrument masks the biological signal of interest. As expected, while residual variance constituted the largest single component, plate_id and peptide intensity made substantial contributions, with the number of quantified precursors per cell contributing a smaller but non-negligible fraction (**Fig. 5j, l**).

After correcting for plate identity, the latent factor representation underwent a striking reorganization, where Astral and timsTOF Ultra2 instruments no longer formed separate clusters, and all nine laboratories became interconnected (**Fig. 5m**). This demonstrates that the cell-line specific differences are conserved after the correction for instrument site and plate-level batch effects. Following plate correction, the two cell types represent distinguishable populations with distinct patterns concerning within-plate spatial effects. Variance partitioning confirmed that residual variance now dominates overwhelmingly, with a subtle contribution from well position within the plate (**Fig. 5m-n**). Correction for both plate identity and within-plate position yielded a fully resolved separation of HEK293T and K562 cells in the latent factor representation (**Fig 5o**). Residual variance accounted for close to 100% of the remaining variance, confirming that the cell-type identity has been successfully isolated from all modeled technical sources (**Fig. 5p**).

Instrument-specific variability of raw protein quantities suppresses cell-type specific differential abundance, which is corrected through batch-correction, directly enabling downstream differential expression analysis (DEA). Taken together, the complete cell type separation after correction provides orthogonal validation that the centralized sample preparation and platform-specific standardized acquisition protocol preserves the biological signal necessary for downstream SCP analysis.

### Cross-laboratory differential expression reveals instrument-dependent biological discovery

Last, we performed DEA using linear mixed models (Msqrob2), with plate and well-location included as covariates (see Methods for detailed description), between HEK293T and K562 cells both within each laboratory independently and in an integrated model combining data across all nine instruments, using FDR < 0.001 and log₂ fold-change > 0.5 as significance thresholds (**Fig. 6a**).

**Figure 6.**
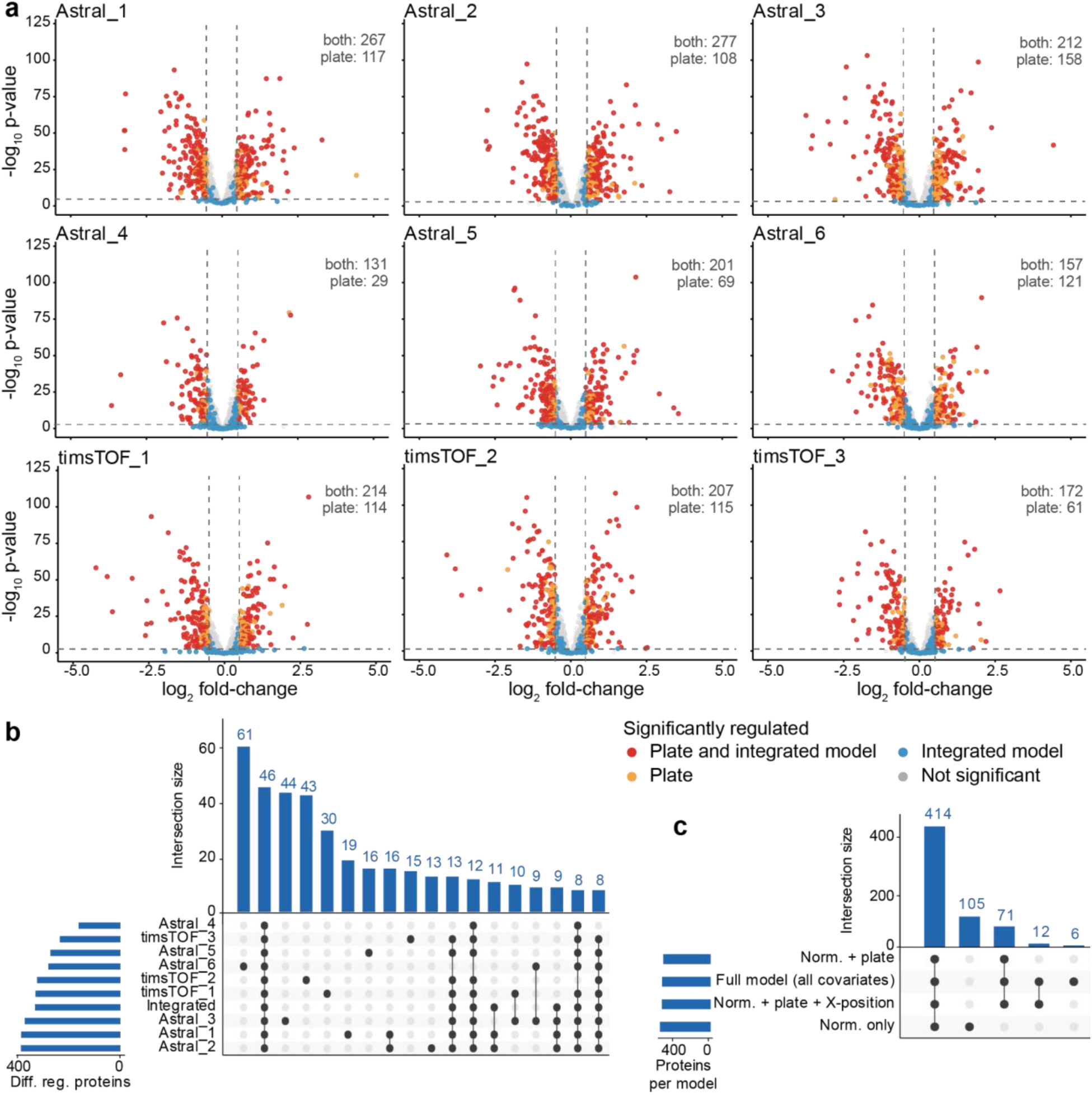
Cross-laboratory differential expression analysis between HEK293T and K562 single cells. **(a)** Per-laboratory volcano plots showing differential protein abundance between HEK293T and K562 single cells (FDR < 0.001, log₂FC > 0.5) for all nine instruments. The number of significant proteins per laboratory is indicated above each plot. Points are colored by their status in the integrated cross-laboratory model: red = significantly regulated in both plate and integrated model, orange = significantly regulated in the plate only, blue = significant in the integrated model only, grey = not significant. Dashed vertical lines indicate log₂FC = 0.5 threshold. UpSet plots showing the overlap of significantly **(b)** regulated proteins across all nine instruments and the integrated model or **(c)** regulated proteins across the models. Bar height indicates the number of proteins in each intersection; the integrated model bar represents proteins identified as significant in the combined cross-laboratory analysis. Horizontal bars on the left hand side indicate the total number of significant proteins per laboratory or in the integrated model.

Per-laboratory DEA revealed substantial variability in the number of significantly differentially abundant proteins recovered across instruments (adjusted p-value < .001; |log2FC| > 0.5), ranging from 160 proteins (Astral_4) to 385 (Astral_2), with most sites identifying between 233 and 328 significant proteins (**Fig. 6a**). This nearly threefold range in per-lab discovery reflects the cumulative effect of instrument-specific differences in identification depth, data completeness, and quantitative precision documented in earlier figures. Notably, Astral_4, the only instrument operated with ABIRD in place of FAIMS, yielded the fewest DE proteins despite demonstrating the tightest QC reproducibility and lowest precursor-level CVs across all instruments (**Fig. 3c**; **Fig. 6a-b**). This suggests that the ABIRD background suppression is beneficial for quantitative precision on clean peptide standards but may alter the sampling of lower-abundance differentially expressed proteins in the more complex single-cell matrix. Highlighting proteins that were significant across the different laboratories, shown in orange (up in K562) and purple (down in K562), reveals a small core of robustly regulated proteins across all laboratories (20 upregulated and 26 downregulated) (**Supplementary Fig. 4**). Most per-lab significant proteins are site-specific discoveries that do not replicate in the integrated analysis and do not correlate with the overall number of identifications (Figure 6a). Laboratories yielding fewer significant DE proteins show a proportionally higher fraction of integrated-model hits among their significant set, suggesting that more stringent per-lab identification thresholds preferentially retain the most reproducible biology.

Examining the overlap of significantly upregulated and downregulated proteins across all nine laboratories using UpSet plots revealed a largely site-specific landscape of DE proteins in both directions (259 upregulated and 255 downregulated; **Fig. 6b-c**). The Astral_6 yielded a disproportionately large number of unique DE proteins (n=61), consistent with its plate being stored for the longest period prior to acquisition, underscoring the importance of monitoring all pre-analytical variables in multi-site or multi-batch studies. The integrated cross-laboratory model recovered 330 DE proteins, representing among the most consistently regulated candidates across the cohort and constituting a reproducibility-filtered reference set for downstream interpretation. The proteins shared across the largest number of individual laboratories, and those recovered by the integrated model, represent the highest-confidence candidates of HEK293T versus K562 biology. Whether the remaining laboratory-unique DE proteins reflect genuine instrument-specific identifications, site specific noise characteristics or sample preparation differences will be subject of future investigation. Taken together, these results demonstrate that while all nine laboratories successfully detect biologically meaningful differences between cell types, the specific proteins identified as differentially abundant are heavily site-dependent when analyzed in isolation. This highlights the need for the community-wide quality control framework described in this study as a prerequisite for meaningful data integration across laboratories and instrument platforms.

## Conclusions

SCP is advancing faster than the quality standards needed to evaluate, compare and validate the data it generates. Yet without such standards, the field risks building its biological conclusions on a foundation that has not been systematically tested. This study, conducted as part of the HUPO SCI, addresses that gap directly by providing the first empirically validated, multi-site quality control framework for SCP. We have systematically deconvolved the contributions of sample preparation, instrument performance, site-specific handling, and computational choice to the total variability observed in SCP data to demonstrate that biologically meaningful cell-type differences are reproducibly detected when appropriate controls are applied.

A central and perhaps initially surprising finding of this study is that software choice is a primary determinant of both identification depth and quantitative accuracy in SCP, exceeding the contribution of instrument identity within each platform. The nearly five-fold range in precursor identifications and substantial differences in quantitative error observed across the six benchmarked tools applied on identical raw data underscores the need for transparent software benchmarking. Indeed, this means that two researchers analyzing the same raw data with different tools may reach different biological conclusions! Peptides identified across all tools provide a high-confidence core identification set that balances the quantitative error and the number of identifications. Based on this, Spectronaut directDIA was selected as the primary analysis tool for this subsequent single-cell processing. Critically, this observation could only be made by inclusion of a dedicated quantitative QC benchmark.

The 384-well plate layout embeds four orthogonal QC tiers alongside 216 single cells per plate, providing the basis to isolate distinct sources of variability. Blank injections confirmed the absence of carryover across all sites and validated the integrity of the sealed-plate shipment workflow. Processing and sorting controls identified cell-leakage background in one plate, which can inflate single-cell identifications. Importantly, these additional identifications were flagged during subsequent processing and DEA. RT stability monitoring enabled the prospective detection and exclusion of injections affected by LC misconfiguration, while plate identification heatmaps localized site-specific QC variability to pipetting errors rather than instrument failure. Each QC tier caught something the others might have missed. Taken together, these four layers of quality control combine chromatographic stability, plate integrity, longitudinal acquisition trends, and data completeness into a deployable, universal QC framework for SCP.

One of the most consequential findings for multi-site study design is that cross-instrument quantitative correlations are impacted by instrument vendors. While the inter-instrument correlations are dominated by instrument vendors, Astral_4 operated with ABIRD in place of FAIMS Pro Duo clusters far from the other Astral instruments in a pair-wise analysis. While Astral_4 produced the tightest CVs and most reproducible identifications from QC standards across all instruments, it yielded the fewest differentially expressed proteins in the single-cell DEA. This suggests that the ion pre-filtering on clean standards may not be identical for the more complex single-cell matrix, and warrants further systematic investigation. As expected, the intra-laboratory correlation resulted in the optimal Pearson correlations for both QC and single-cell samples (∼0.9) immediately followed by the tims-tims correlations (0.8). Interestingly, the Astral-Astral correlations decreased to 0.7, which was slightly recovered by removing Astral_4 from the analysis (0.7-0.79). The largest spread of correlation was observed in tims-Astral comparisons, even though some individual instruments achieved a correlation of up to 0.75 in single-cell samples across vendors. The conclusion here is fundamental: data integration across sites requires explicit batch correction strategies irrespective of instrument choice. Sequential correction for plate identity and within-plate position using linear models resolved the position-driven clustering which dominated the uncorrected data and yielded clean separation of HEK293T and K562 cells. Equally important, the recovery of consistent, biologically interpretable cell-type differences after correction provides orthogonal validation that the centralized sample preparation protocol preserves the biological signal necessary for SCP-based discovery.

Cross-laboratory differential expression analysis revealed that the specific proteins identified as significantly regulated between HEK293T and K562 cells are heavily site-dependent when laboratories are analyzed in isolation. Per-site discovery ranged up to 385 significant proteins, with the majority being laboratory-unique and not recovered by other sites. The regulated proteins identified by the integrated multi-site model yielded the most consistent and reproducibility-filtered set of DE candidates across the cohort. While the multi-site model represents low significant hits in the individual sites, future studies will demonstrate the biological impact of robustly regulated DEs and reinforces the need for QC and normalization frameworks described here as a prerequisite for meaningful cross-site data integration.

To facilitate adoption of these standards by the broader SCP community, this study provides: (i) a fully annotated raw and processed benchmarking dataset deposited on Zenodo and Mass Dynamics; (ii) software benchmarking results applying developer-recommended settings across six tools on identical input data; (iii) instrument vendor-specific and platform-comparative data evaluations; (iv) intra- and inter-laboratory variance estimations across two cell types and four QC sample categories; and (v) a multi-tier, plate-embedded QC framework with explicit design specifications, injection sequences, and post-acquisition analysis recommendations.

The field of SCP has seen tremendous innovation at instrument and software level but what has been missing is the shared quality and benchmarking infrastructure that allows to build on each others work. As SCP transitions from methodological innovation to biological and clinical application, the reproducibility, comparability, and interpretability of the data it generates will be determined not by any single technological advance, but by the collective quality of the experimental and analytical standards the community chooses to adopt. We believe this study provides critical elements to the empirical groundwork for such standards.

## Acknowledgments and funding

L.M. acknowledges funding from the Horizon Europe Projects BAXERNA 2.0 [101080544] and COMBINE [101191739], and from the Ghent University Concerted Research Action [BOF21/GOA/033]. L.M. and C.Ca. are further supported by the CHIST-ERA project ODEEP-EU [G0GDV23N] and FWO SRN [W005325N]. S.v.P., T.C and L.M. acknowledge funding from the Research Foundation Flanders (FWO) [1SA7M26N, 12A8W25N, G010023N, G028821N]. C.Ca. acknowledges the Agence Nationale de la Recherche via the French Proteomic Infrastructure (ProFI UAR2048; ANR-10-INBS-08-03 and ANR-24-INBS-0015), the Région Grand-Est (SC-Proteomics and Proteomique projects), the Eurométropole de Strasbourg, the ITMO Cancer of Aviesan within the framework of the 2021-2030 Cancer Control Strategy, on funds administered by Inserm (ProteomiSC project), for equipment funds. J.B.R., C.K. and C.Ca. acknowledge the Interdisciplinary Thematic Institute IMS, the drug discovery and development institute, as part of the ITI 2021-2028 program of the University of Strasbourg and CNRS, supported by IdEx Unistra (ANR-10-IDEX-0002), and by SFRI-STRAT’US project (ANR-20-SFRI-0012).

## Materials and Methods

### Centralized Sample Preparation

HEK293T and K562 cells were cultured under standard conditions in Dulbecco’s Modified Eagle’s Medium supplemented with 10% Fetal Bovine Serum and Penicillin-Streptomyocin at 37°C and 5% CO_2_. Cells harvested with Gibco Trypsin-EDTA and washed 3 times (300 rpm for 4 min) immediately prior to plate preparation. Single cells were isolated and deposited using a cellenONE X1 Neo (Cellenion, Lyon, France), according to the following parameters: minimum 19 µm and maximum diameter 26 µm with 1.65 elongation.

Each wells rows C-N and columns 2-23 of the 384-well plate received 1 μL of digestion master mix consisting of 100 mM triethylammonium bicarbonate (TEAB), 0.2% n-dodecyl-β-D-maltoside (DDM), and 5 ng Trypsin/Lys-C Mix, Mass Spec Grade (Promega, V5071). Individual cells were subsequently dispensed into their respective wells by the cellenONE where HEK293T cells were deposited in rows C-N and columns 4-12 and K562 in rows C-N and columns 13-21. Sorting controls (rows C-N and columns 22-23) received one droplet of the sorting liquid without a cell (negative isolation on the cellenONE). Plates were then incubated at 45°C for 2 h with continuous rehydration cycles (500 nL per well rows C-N columns 2-23) performed by the cellenONE to counteract evaporation. Digestion was quenched by addition of 1 μL of 0.1% formic acid (FA) containing 1% dimethyl sulfoxide (DMSO). 2 µL of quantitative QC mix consisted of a three-proteome mixture of human, yeast (*Saccharomyces cerevisiae*), and *E. coli* tryptic digests prepared at two distinct abundance ratios (ratio A: 15% yeast / 20% *E. coli*; ratio B: 30% yeast / 5% *E. coli*; human proteome constituting the remainder in both cases), each injected at a total protein amount of 200 pg was added to row B 2-12 and row O 13-23. Qualitative sensitivity controls consisted of 2 µL Pierce HeLa Protein Digest Standard (Thermo Fisher Scientific, #88328) at 200 pg per injection were added to row B 13-24 and row O 2-12. Blank wells received 2 µL quenching reagent (0.1% FA and 1% DMSO) in row C-N and columns 1 and 24. Plates were sealed with aluminum foil, briefly centrifuged and stored at −80°C until shipment on dry ice to participating laboratories.

Upon receipt, participating laboratories thawed plates at room temperature for 10 min, followed by brief centrifugation at 200*g* for 1 min. Each site then appended site-specific qualitative controls (Pierce HeLa Protein Digest Standard prepared locally using the same dilution protocol, 200 pg per injection in row A column 1-12, row B column 1, row O column 24, row P column 13-24) and site-specific blank injections (0.1% FA; row A column 13-24, row B column 24, row O column 1, row P column 1-12) to monitor for any signal variation introduced by shipment or freeze-thaw cycling. All injections across the complete plate were acquired in a randomized block order as indicated in the supplemented metadata files, alternating blocks of QC injections with blocks of single-cell injections. The final injection volume per plate was determined in well C3 prior to injection due to possible fluctuations during the rehydration procedure. Plates were then sealed with silicone sealing mats for injection (Axygen™ AxyMats™ Sealing mats #14-222-017).

### Thermo Fisher Vanquish Neo setup

Chromatographic separation was carried out on a Vanquish Neo UHPLC system (Thermo Fisher Scientific) using a two-column setup: a PepMap C18 trapping column (5 mm × 300 μm, Thermo Fisher Scientific) for online desalting, coupled to an Ionopticks Aurora Series C18 analytical column (8 cm × 75 μm).

The chromatographic gradient was initiated at 4% B (20% H_2_O, 79.9% acetonitrile, 0.1% FA) and a flow rate of 800 nL, ramped to 12% B within 0.075 min, followed by 18% within 0.188 min, 35% in 0.8 min and 99% within 0.05 min. Held at 99% for 0.338 min, flow rate was decreased to 100 nL and held for 6.249 min afterwards increased to 800 nL and 4% B in 0.05 min. For the indicated laboratories using 0.1% FA in 100% acetonitrile the %B was adapted as follows: 3.2% at 800 nL during gradient initiation, ramped to 9.6% in 0.075 min, to 14.4% in 0.188 min, 28% in 0.8 min, to 80% in 0.05 min held for 0.338 min. Flow rate was decreased to 100 nL in 0.05 min held for 6.249 min and increased to 0.8 with 3.2%B within 0.050 min.

### timsTOF Ultra2 Sample Acquisition

Acquisition on the timsTOF Ultra2 was performed in diaPASEF mode accumulating a ramping from 0.7-1.3 1/*K*_0_ in 100 ms with High Sensitivity Mode enabled and a collision energy slope of 0.6-1.6 starting for the IM range of 20-59. DIA PASEF windows were dynamically placed from 400-1000 *m/z* as follows.

**Table 1:**

| Cycle Id | Start IM [$1/K_0$] | End IM [$1/K_0$] | Start Mass [$m/z$] | End Mass [$m/z$] |
| --- | --- | --- | --- | --- |
| 1 | 0.97 | 1.45 | 722.28 | 767.38 |
| 1 | 0.84 | 0.97 | 557.33 | 587.32 |
| 1 | 0.64 | 0.84 | 400 | 434.58 |
| 2 | 0.99 | 1.45 | 766.38 | 818.9 |
| 2 | 0.89 | 0.99 | 586.32 | 617.81 |
| 2 | 0.64 | 0.89 | 433.58 | 470.59 |
| 3 | 1.01 | 1.45 | 817.9 | 882.96 |
| 3 | 0.905 | 1.01 | 616.81 | 650.32 |
| 3 | 0.64 | 0.905 | 469.59 | 501.26 |
| 4 | 1.045 | 1.45 | 881.96 | 956.47 |
| 4 | 0.925 | 1.045 | 649.32 | 684.39 |
| 4 | 0.64 | 0.925 | 500.26 | 529.81 |
| 5 | 1.11 | 1.45 | 955.47 | 1000 |
| 5 | 0.955 | 1.11 | 683.39 | 723.28 |
| 5 | 0.64 | 0.955 | 528.81 | 558.33 |

### Orbitrap Astral Sample Acquisition

Acquisition on the Orbitrap Astral was performed for 7.8 min in Peptide mode. The spray voltage was 1.9 kV with 240°C ion transfer tube temperature, using standard resolution on FAIMS with 3.6 L/min carrier gas flow at CV of -48 V. Full Scans were acquired and 240,000 resolution across 400-800 *m/z* scan range with an RF lens of 45%, 500% AGC target with an absolute AGC target of 5×10^6^ for maximum 100 ms injection time. DIA windows were dynamically placed starting from 400-800 *m/z* (400-437, 437-454, 454-472, 472-485, 485-498, 498-511, 511-523, 523-536, 536-547, 547-599, 599-572, 572-584, 584-596, 596-611, 611-626, 626-645, 645-664, 664-688, 688-722, 722-800 *m/z*). HCD collision energy was set to 25%, ions were collected until an AGC target of 500% (absolute value of 5×10^4^) or maximum injection time of 40 ms was reached. One indicated laboratory instead of FAIMS utilized Active Background Ion Reduction Device (ABIRD, ESI Source Solutions, Woburn, MA) for ion pre-filtering.

### Protein sequence databases

Two databases were used for peptide and protein identification. The three-species benchmark mixture was searched against a combined database of 20,405 human (Homo sapiens, OX=9606), 4,402 Escherichia coli K-12 (OX=83333), and 6,066 Saccharomyces cerevisiae (OX=559292) canonical protein sequences, supplemented with 381 common contaminant sequences (31,254 entries total). Human and microbial sequences were downloaded from UniProt on 17 October 2025 (reviewed, canonical isoforms). Contaminant sequences were obtained from the Universal Protein Contaminant Library for DDA and DIA proteomics (HaoGroup-ProtContLib, GitHub); contaminant entries carry a “Cont_” prefix in the accession field. Single-cell human samples were searched against a human-only database of the same 20,405 canonical human sequences supplemented with the same 381 contaminant sequences (20,786 entries total).

### DIA search parameters

#### Species-mixture benchmarking

Raw DIA files of the species-mixture samples were processed independently per plate identifier using DIA-NN (2.6.0), Spectronaut (20.5), AlphaDIA (2.0.1), CHIMERYS (5.0.0), FragPipe (24.0, MSFragger 4.4.1, quantification via DIA-NN 2.2.0), and PEAKS Online (13). All searches used identical FASTA databases and were performed at 1% FDR at both precursor and protein level. Each tool was configured according to its developer-recommended workflow for library-free DIA analysis, where parameters were kept as default as best as possible. Key parameters are summarised in Table 2.

**Table 2:**
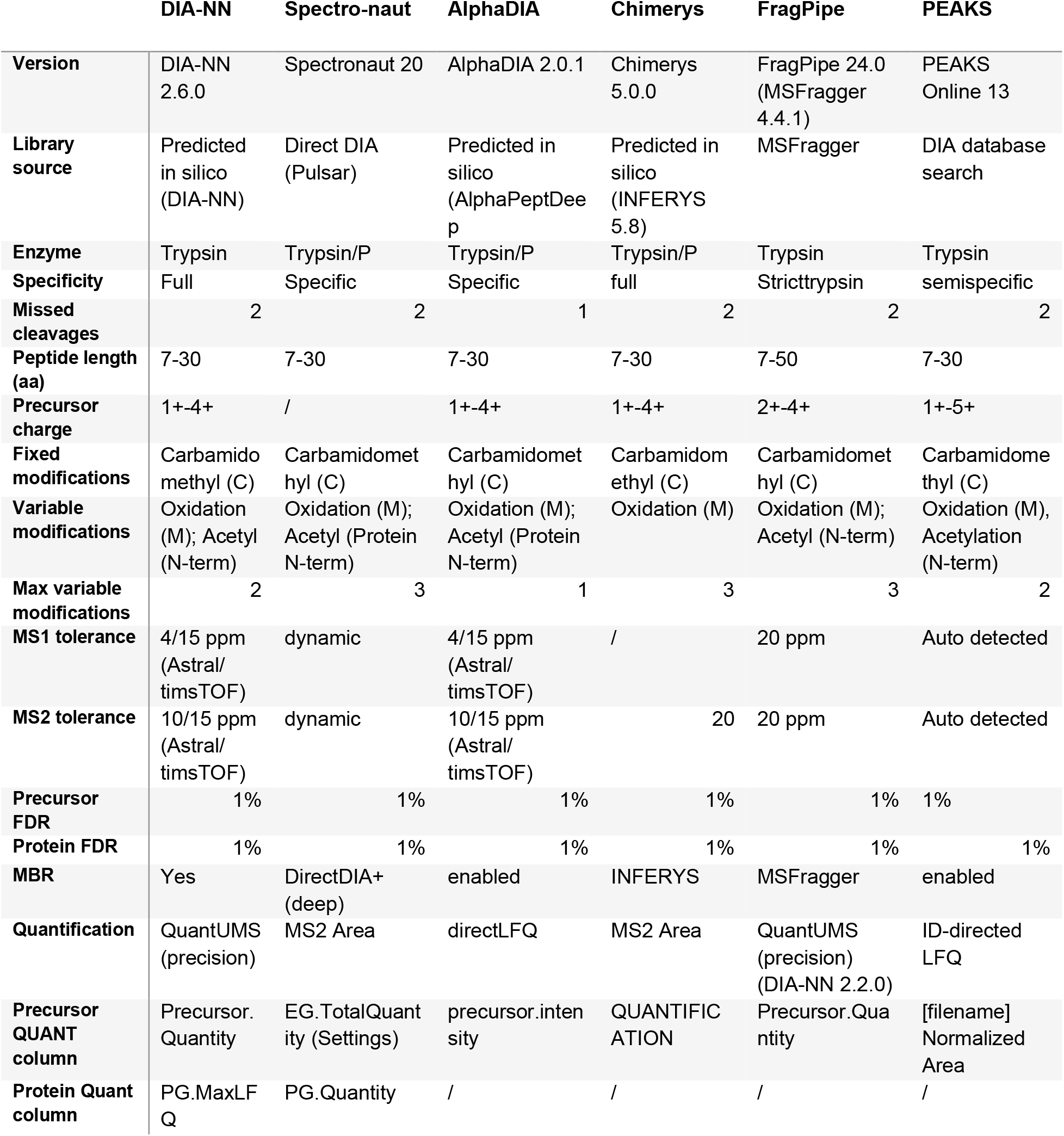

Library generation differed by tool: DIA-NN and AlphaDIA generated predicted in silico libraries using DIA-NN and AlphaPeptDeep, respectively; CHIMERYS used INFERYS 5.8; Spectronaut performed directDIA analysis via its Pulsar engine; FragPipe used a pseudo-DDA strategy via MSFragger; and PEAKS Online applied a DIA database search. MS1 and MS2 tolerances were set to 4 and 10 ppm (Astral) or 15 ppm in both dimensions (timsTOF Ultra2) for DIA-NN and AlphaDIA, 20 ppm in both dimensions for FragPipe, and 20 ppm (initial fragment tolerance) for CHIMERYS; Spectronaut and PEAKS determined tolerances automatically. PEAKS Online applied a score-based precursor filter (−10lgP ≥ 20), corresponding to approximately 1% FDR as defined by the vendor. Precursor-level quantification used QuantUMS precision mode (DIA-NN, FragPipe), MS2 area (Spectronaut, CHIMERYS), directLFQ (AlphaDIA), and ID-directed LFQ (PEAKS). For CHIMERYS and PEAKS, internal run-wise normalization was unavailable or disabled; a systematic offset in their log2 fold-change distributions was corrected by post-hoc TIC normalization (see Quantitative accuracy and software benchmarking).

#### Complete single-cell dataset

Spectronaut 20 was used in directDIA mode for the full dataset. Identification was performed by the Pulsar engine using the directDIA+ (Deep) workflow against the human database described above. Trypsin/P was set as the cleavage enzyme with full specificity and a maximum of two missed cleavages, and peptide length was restricted to 7-30 amino acids. Carbamidomethylation of cysteine was a fixed modification; N-terminal protein acetylation and methionine oxidation were variable modifications (maximum three per peptide). N-terminal methionine cleavage was enabled. PSM, peptide, and protein group FDR were each set to 1%, and MS1 and MS2 tolerances used dynamic calibration. Decoys were generated internally by mutation using a dynamic fraction of 10% of the library size, with neural network-predicted fragment ions as the preferred fragment source.

Per-plate SNE result files were combined using the Spectronaut command-line combine function (SNE-merge), which re-ran all downstream DIA steps jointly across plates: protein inference (IDPicker), CScore normalization, interference correction (minimum two MS1 and three MS2 ions), global precursor FDR estimation, experiment- and run-level protein FDR estimation, cross-run quantity normalization (automatic strategy), and background noise subtraction. Precursor Q-value cutoffs were 1% at both run and experiment level; protein Q-value cutoffs were 1% at experiment level and 5% at run level. Quantification was based on MS2 fragment ion peak area using the top one to three peptides per protein group (mean precursor area per peptide, mean peptide quantity per protein). No missing-value imputation was performed. This dataset was used for all downstream analyses.

The pipeline was executed in parallel using DIA-NN (2.6.0), reproducing the same figure panels with different search toll output. Each plate was searched independently against an in silico predicted spectral library generated with its deep-learning predictor:

Command:

> *diann-linux --predictor --fasta human_contaminant_revi_2025_10_17.fasta --fasta-search --met-excision --min-pep-len 7 --max-pep-len 30 --min-pr-mz 300 --max-pr-mz 1800 --min-pr-charge 1 --max-pr-charge 4 --min-fr-mz 200 --max-fr-mz 1800 --cut K*,R* --missed-cleavages 2 --unimod4 --var-mods 2 --var-mod UniMod:35,15.994915,M --var-mod UniMod:1,42.010565,*n --peptidoforms --no-prot-inf*

Plates were then searched against this library using a two-pass approach with match-between-runs: the first pass generated an empirical spectral library from the DIA data with ID, RT, and IM profiling, and the second pass reanalyzed the experiment with this empirical library. MS2 and MS1 tolerances were 10 and 4 ppm for Astral data and 15 ppm for both levels for timsTOF data, with a scan window of 8. Protein grouping used UniProt protein name (--pg-level 1). MBR output tables (report.parquet) from all plates were concatenated and filtered to FDR levels equivalent to Spectronaut: run-level precursor Q-value (Q.Value) < 0.01, run-level protein group Q-value (PG.Q.Value) < 0.05, global peptidoform Q-value (Global.Peptidoform>Q.Value) < 0.01, and global protein group Q-value (Global.PG.Q.Value) < 0.01.

### Metadata processing

One SDRF file was provided per plate, containing acquisition and sample metadata and well positions. cellenONE reports generated during dispensing contained morphological measurements, dispensing QC flags, and paths to brightfield images. These two sources were joined on well position and plate number. Manual per-well QC annotations derived from plate preparation notes were applied as four boolean flags: Doublet (two co-dispensed cells), Dividing (cell in mitosis), Dying (morphological signs of cell death), and CellInBlank (cell accidentally dispensed into a designated blank well).

The merged table was processed as follows: sample names were homogenized across plates; timestamp columns were parsed to datetime components; injection order was normalized between 0 and 1 based on gradient-time timestamps; and missing values were imputed with sentinel values (-1 for numeric fields; “not applicable” or “not available” for text fields). Three sites required additional corrections: Astral_4 and timsTOF_2 performed reinjections of failed acquisitions, which were annotated in the metadata; Astral_6 lacked gradient-time annotations, which were copied from Astral_3 (injection order was consistent by design); and timsTOF_3 had transposed sample-type labels for site-specific QC and blank wells, which were corrected. The final metadata table was saved as a single parquet file and added to the AnnData object for all downstream analyses.

### AnnData Object construction

Search results from Spectronaut and DIA-NN (2.6.0) were converted into AnnData objects. For each search engine, four objects were constructed: precursor- and protein-level quantification for human samples, and the same two levels for the quantitative QC mixture samples (Table 3 shows which columns were used for abundance information and precursor identifier). Modification notation was standardized to UNIMOD accession format for both engines. Results were joined to the metadata table on plate identifier and sample name, and the merged long-format table was pivoted into a sparse samples × features intensity matrix. Additional layers stored per-precursor quality scores, RT, and, for DIA-NN, IM values. A log-transformed intensity layer was added to all objects, and proteins with a “Cont_” prefix were flagged as contaminants in the feature annotations. Objects were written to disk in .h5ad format and are available on Zenodo.

**Table 3:**

|  | **Spectronaut** | **DIA-NN** |
| --- | --- | --- |
| **Precursor ID** | FG.PrecursorIon | Modified.Sequence/{Precursor.Charge} |
| **Precursor Quant** | EG.TotalQuantity | Precursor.Quantity |
| **Protein ID** | PG.ProteinGroups | Protein.Group |
| **Protein Quant** | PG.Quantity | PG.MaxLFQ |

### Quantitative accuracy evaluation

For the three-species QC analysis, a precursor was required to be detected in at least three raw files within each of the two mixing conditions (A and B), and precursors mapping to more than one species were removed. Intensities were log2-transformed, averaged within each condition, and the measured fold-change taken as the difference of condition means (log2_A_vs_B = log_intensity_mean_A − log_intensity_mean_B). The signed deviation from the expected value was ε = log2_A_vs_B − log2_expected_ratio, with expected per-species ratios of 0, -1, and 2 for human, yeast, and E. coli, respectively. The mean absolute difference between measured and expected fold-changes was the average |ε| across all precursors passing the detection filter. For CHIMERYS and PEAKS, TIC normalization was applied before fold-change calculation by dividing each run’s quantity by its summed signal (CHIMERYS: sum of the QUANTIFICATION column; PEAKS: sum of the Normalized Area column per sample).

Fold-change distributions were visualized per laboratory as Gaussian kernel-density estimates with expected ratios drawn as vertical reference lines. For MA-plots, mean abundance was defined as 0.5 × (log_intensity_mean_A + log_intensity_mean_B).

For UpSet plots, precursor identity overlap was computed across all six workflows after applying the per-condition detection filter (minimum three identified precursors in each of conditions A and B). Each tool’s identified set was defined as the set of precursor ion strings (stripped sequence with modifications plus charge, e.g. AAAAM[UNIMOD:35]AAALQAK/2). The intersection was the set of precursors identified by all tools; tool-unique sets were precursors identified by exactly one tool. Quantitative accuracy across these subsets was compared using absolute fold-change errors (|ε|). Physicochemical properties were derived directly from each precursor ion string: peptide length, precursor charge, missed cleavages (count of internal K or R residues not followed by proline), and modification status (unmodified, oxidation, acetylation, or acetylation plus oxidation). Values were computed both for all identified precursors per tool and for the subset excluding the all-tools intersection, with laboratories grouped by platform (Astral, timsTOF).

### QC metrics and analysis

#### Identification heatmap

Detection was visualized as a sample-by-feature heatmap, where features were ordered by identification frequency in the single-cell samples, and samples were grouped by condition, with the single-cell conditions randomly down-sampled to at most 50 samples per condition for visualization purposes.

#### Coefficient of variation

The coefficient of variation was used to summarise quantitative reproducibility within each laboratory and condition. Contaminant features were removed before the calculation and the calculation was run for the four conditions (K562, HEK293T, Site specific QC and QC - Hela) at both the precursor and the protein level.

Features were grouped by the pair (plate_idx, source name), that is, one group per laboratory and condition. For each feature within a group the percentage coefficient of variation was computed on raw (linear) intensities from adata.X as

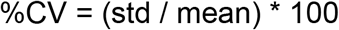

with the standard deviation computed using the N-1 denominator (ddof = 1) and with NaN-aware mean and standard deviation. A feature required more than one valid (non-NaN) observation within a group to yield a value.

For the plotted distributions, a feature was required to have at least ten valid observations within the group (min_n = 10). Two feature sets were reported per condition: (i) the full set of quantified features, and (ii) a “common proteome”-set where features for a condition should be quantified at least 10 samples on each plate. Both figures are on GitHub. In the manuscript, the full feature set version is shown.

#### Completeness

Data completeness was quantified over a grid of thresholds between 0 to 1 in steps of 0.01. Two complementary curves were computed per laboratory (colored line) and pooled across laboratories (dotted, black line), at both precursor and protein level:

- Feature-level completeness: The fraction of samples in which a feature was detected. The feature-retention curve is the percentage of features detected in at least *t* percent of samples.
- Sample-level completeness: The fraction of detectable features observed in a sample. The sample-retention curve is the percentage of samples covering at least t percent of features.

Curves were computed separately for the single-cell conditions (K562, HEK293T) and the QC conditions (QC - Hela, site specific QC).

Per-sample completeness was stored for the single-cell (SC) and QC (QC) condition sets, which expresses each sample’s completeness relative to the number of features detectable for that condition within the plate. A sample was flagged as ‘completeness outlier’ when its per-sample precursor completeness is below 0.25 in the single-cell set or in the QC set.

The 0.25 threshold was chosen by inspection of the completeness curves. The flag was computed on the precursor object and propagated to the protein object. Samples flagged as completeness outliers were excluded from the RT, injection order, correlation and other downstream analyses.

#### Injection order analysis

The per-plate injection order was derived from the acquisition timepoints. Within each plate, the timepoints were ranked and the ranks were rescaled between 0 and 1. The identification counts were expressed as a percentage of the per-condition, per-day maximum. For each plate and condition, an ordinary least-squared regression was performed on the normalized timepoints and identification count percentages.

#### Chromatographic stability

Chromatographic reproducibility was assessed from precursor apex RT, taken from the Spectronaut column EG.ApexRT. These were first normalized within each sample to the interval [0, 1] using the earliest and latest eluting precursor in that sample. For each sample, the normalized RTs were binned into 30 equal-width bins. For each sample, the proportion of precursor identifications in each bin was plotted at the midpoint, giving one RT distribution per sample. These distributions were computed per plate for the four conditions: K562, HEK293T, Site specific QC and QC - Hela.

### Correlation analysis

#### Rationale

Pairwise sample correlations were used to quantify quantitative reproducibility within and between laboratories and instrument platforms. Because missingness differs systematically between laboratories, a naive correlation would be confounded by the number and identity of co-observed features. To make comparisons fair, a reference protein set with equalised missingness across laboratories was constructed before correlations were computed.

#### Reference-set construction

The reference set was built at the protein level on the log2 for the four conditions: K562, HEK293T, Site specific QC, and QC - Hela. Reference-set construction per condition was performed in three steps:

1. **Sample filtering:** Samples with completeness values below 0.25 were filtered. See section on completeness.
2. **Reference protein selection:** A protein was retained only if it was detected in at least 75% of samples in every plate.
3. **Missingness equalisation:** For each retained protein, the target missingness was set to the maximum (worst-case) per-plate missingness across all plates. In every plate, observed values were then randomly masked until that plate’s per-protein missingness matched the target. This ensures that all plates contribute the same fraction of missing values per protein, so that cross-laboratory correlations are not driven by differences in completeness.

This filtering resulted in 758 (K562), 735 (HEK293T), 1376 (Site specific QC), and 1388 (QC - Hela) reference proteins.

#### Correlation calculation

Within each condition, every sample pair was correlated over the proteins co-observed in both samples without imputation. Both Pearson (shown in the main figures) and Spearman correlations (available on GitHub) were computed. A sample pair was required to share at least 100 co-observed proteins. Although the overlap between each sample pair is different, the reference-set construction makes this effect minimal while keeping enough proteins to perform a correlation analysis.

### Variance partitioning and batch effect correction

Variance partitioning quantified how much of the per-precursor variability in the single-cell data was explained by technical covariates (normalisation proxies, plate, within-plate position, injection order) versus biology (cell type), and how the variance structure and the low-dimensional embedding changed as technical effects were progressively removed. The analysis was performed at the precursor level with scplainer from the scp package (version 1.20.0).

#### Input and cell selection

Modelling used the precursor-level AnnData object with the X_log2-layer as the precursor intensity matrix. Only the single-cell conditions were used where single-cells were filtered based on the completeness filter (described above) and precursors from contaminant proteins were filtered out. After filtering, the matrix was converted into a Qfeatures object (available on Zenodo). Two per-cell normalisation proxies were used: medianIntensity Peptide, the per-cell median log2 precursor intensity, and n_precursors, the number of precursors quantified per cell. A per-cell median coefficient of variation (medianCVperCell, grouped by Protein.Group, n = 5) was computed for quality inspection only and was unused otherwise.

#### Model and covariates

A separate weighted linear model was fitted per precursor with the scpModelWorkflow. The full model formula:

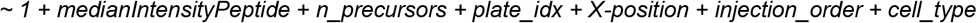

with medianIntensityPeptide and n_precursor as normalisation covariates, plate_idx (plate identifier), X-position (within-plate column position), and injection_order as batch covariates; and cell_type as the biological covariate.

Batch effects were removed sequentially with scpRemoveBatchEffect (intercept = True). Starting from the full fitted model, each stage regressed out a progressively larger set of effects and refitted the reduced model, so that each stage shows the variance explained on top of the effects already removed, which can be visualized in a lower-dimensional space.

1. **Full model**: all covariates above.
2. **Post-normalisation**: remove medianIntensityPeptide and n_precursors; refit ∼ 1 + plate_idx + X-position + cell_type.
3. **Post-plate**: additionally remove plate_idx; refit ∼ 1 + X-position + cell_type.
4. **Post-position**: additionally remove X-position and injection_order; refit ∼ 1 + cell_type, leaving only the biological effect.
5. **Residuals**: remove all of the above and cell_type, giving the unexplained residual matrix with no modelled effect retained.

#### Feature filtering

Because the model is fitted feature-wise, reliability depends on the ratio of observations to parameters (n/p). Precursors with an n/p ratio at or below 20 in the full model were excluded (scpModelFilterNPRatio > 20). The threshold was applied to the full model only, which is the most parameter-rich and therefore most stringent stage, so that features passing there are reliably fitted at every simpler stage. Downstream variance analysis and dimensionality reduction used this selection of precursors, namely 23,037 precursors.

#### Variance analysis

Per-covariate variance was extracted per stage with scpVarianceAnalysis, which returns the percentage of variance explained by each covariate for each precursor. Two views were reported per correction stage: a bar chart of the mean percentage of variance explained per covariate (scpVariancePlot), and violin plots of the per-precursor distribution of variance fractions, ordered by median. Covariates were grouped by functional role (normalisation, plate position, biology) with the residual fraction shown separately.

#### Dimensionality reduction

Low-dimensional structure was obtained by generalized matrix factorization with omicsGMF (version 1.0.0, runGMF), using a Gaussian family. The number of components was set to 4, chosen from the reconstruction-error curve of a rank scan over ranks 1 to 10. A 4-component GMF embedding was computed for each correction stage on common_features.

### Differential expression analysis

Differential protein abundance between the two single-cell types (K562 and HEK293T) was tested with msqrob2, a per-protein ridge-penalised robust mixed-effects model fitted on precursor intensities aggregated to proteins. The analysis was run at two levels: an integrated model across all plates, and an independent per-plate model that treats each plate as a single-laboratory experiment.

The precursor-level AnnData object was loaded and the log2 layer was used as the intensity matrix. Cells were restricted to the two single-cell conditions that were not flagged as completeness outliers (see above), and precursors were kept if observed in at least one retained cell. Precursors were assembled into a QFeatures object. A precursor was retained if it carried a valid, non-contaminant protein-group annotation and belonged to a protein group containing at least one precursor. Precursors were then aggregated to proteins using median polish. Two per-cell covariates were derived, medianIntensityPeptide (per-cell median protein intensity) and n_proteins (number of quantified proteins per cell).

#### Msqrob2 model and testing

Each model was fitted with msqrob2 on the proteins median polish assay using ridge penalisation and robust estimation. The hypothesis test was performed on the null hypothesis that K562 – HEK293T = 0, applying Benjamini-Hochberg method for multiple testing correction. A protein was called significant when its adjusted p-value was below

### 0.001 and the absolute log2 fold-change exceeded 0.5

The integrated analysis fitted four models with a progressively richer set of technical covariates:

Norm: ∼ 0 + cell_type + medianIntensityProtein + n_proteins
Plate: Adds a plate random intercept (1| plate_idx)
Position: Adds a per-plate random slope on column position (0 + X-position | plate_idx)
All: Adds injection_order as a fixed_effect

Note that the plate random intercept and per-plate position slope were entered as uncorrelated terms (1 | plate_id) and (0 + X-position | plate_id). The first term gives each plate its own baseline, absorbing plate-to-plate shifts in overall log2 abundance; the second gives each plate its own gradient across the plate columns (X position), absorbing plate-specific position trends.

Per model, the number of tested and significant proteins were displayed as an UpSet plot representing the overlap between the four models’ significant-protein sets.

#### Per plate analysis

Each plate was analyzed independently to estimate how many integrated hits a single laboratory would recover. Per plate, precursors were aggregated to proteins by median polish and fitted with the same ridge robust linear-mixed model but simplified to:

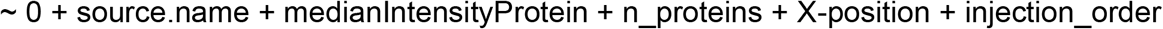

The overlap between the integrated full model and the per-plate results was assessed in two ways. First, on all tested proteins (figures in GitHub). Second, on a commonly quantified protein set, to remove the effect of differing identification sets on the overlap. The models were not refit, but a completeness filter was applied post hoc to the results retaining only proteins detected in at least 25% of cells in at least 5 plates.

## Data and code availability

The raw mass spectrometry data and search results have been deposited in the ProteomeXchange Consortium via the PRIDE repository and will be made available upon publication. AnnData objects, output files from scplainer, MSqRob, and benchmarking results, associated metadata, and brightfield images of isolated single cells are available on Zenodo at 10.5281/zenodo.21181080. Analysis notebooks, figure generation scripts, and supplementary figures based on DIA-NN results (the manuscript presents results obtained with Spectronaut) are available on GitHub at https://github.com/CompOmics/SCI.

## Extended Data

**Extended Figure 1:**
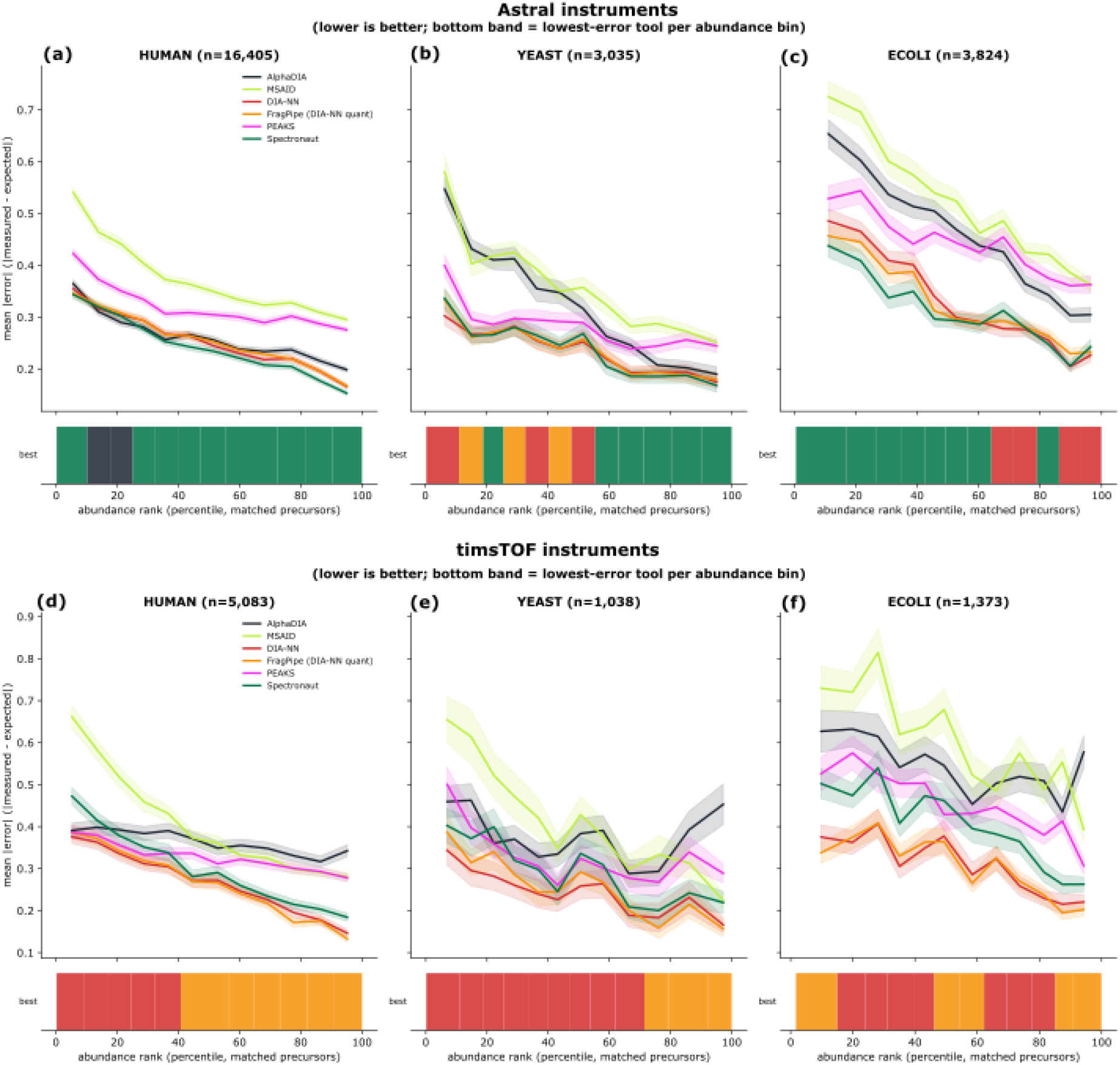
Mean absolute error between measured and expected precursor quantities versus global abundance rank percentile for six DIA software tools (AlphaDIA, MSAID, DIA-NN, FragPipe DIA-NN quant, PEAKS, Spectronaut; colored lines with shaded 95% confidence intervals) across the *quantitative QC* (human - left, yeast - middle, E. coli - right). Results are shown separately for Astral instruments (top) and timsTOF instruments (bottom). The colored bar at the bottom of each panel indicates the software tool with the lowest quantitative error in each abundance bin.

**Extended Figure 2:**
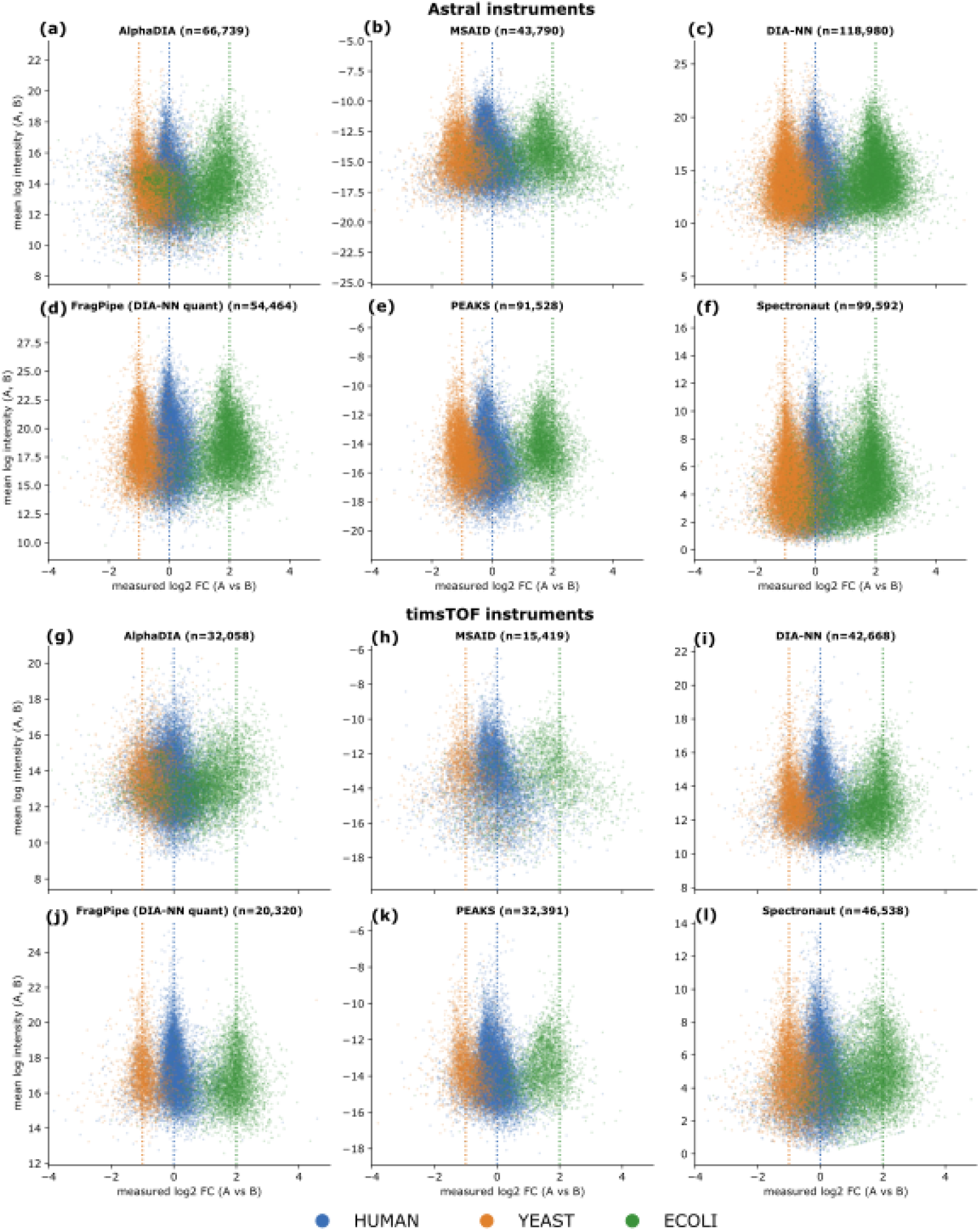
MA plots indicating mean log2 intensity versus log2 fold-change for all matched precursors identified by all of the six DIA software tools: AlphaDIA (**a, g**), MSAID (**b, h**), DIA-NN (**c, i**), FragPipe DIA-NN quant (**d, j**), PEAKS (**e, k**), and Spectronaut (**f, l**), applied to the *quantitative QC*. Results are shown separately for Astral instruments (**a–f**) and timsTOF instruments (**g–l**), colored by species origin. Vertical dashed lines indicate the expected log2 fold-change for each species.

**Extended Figure 3:**
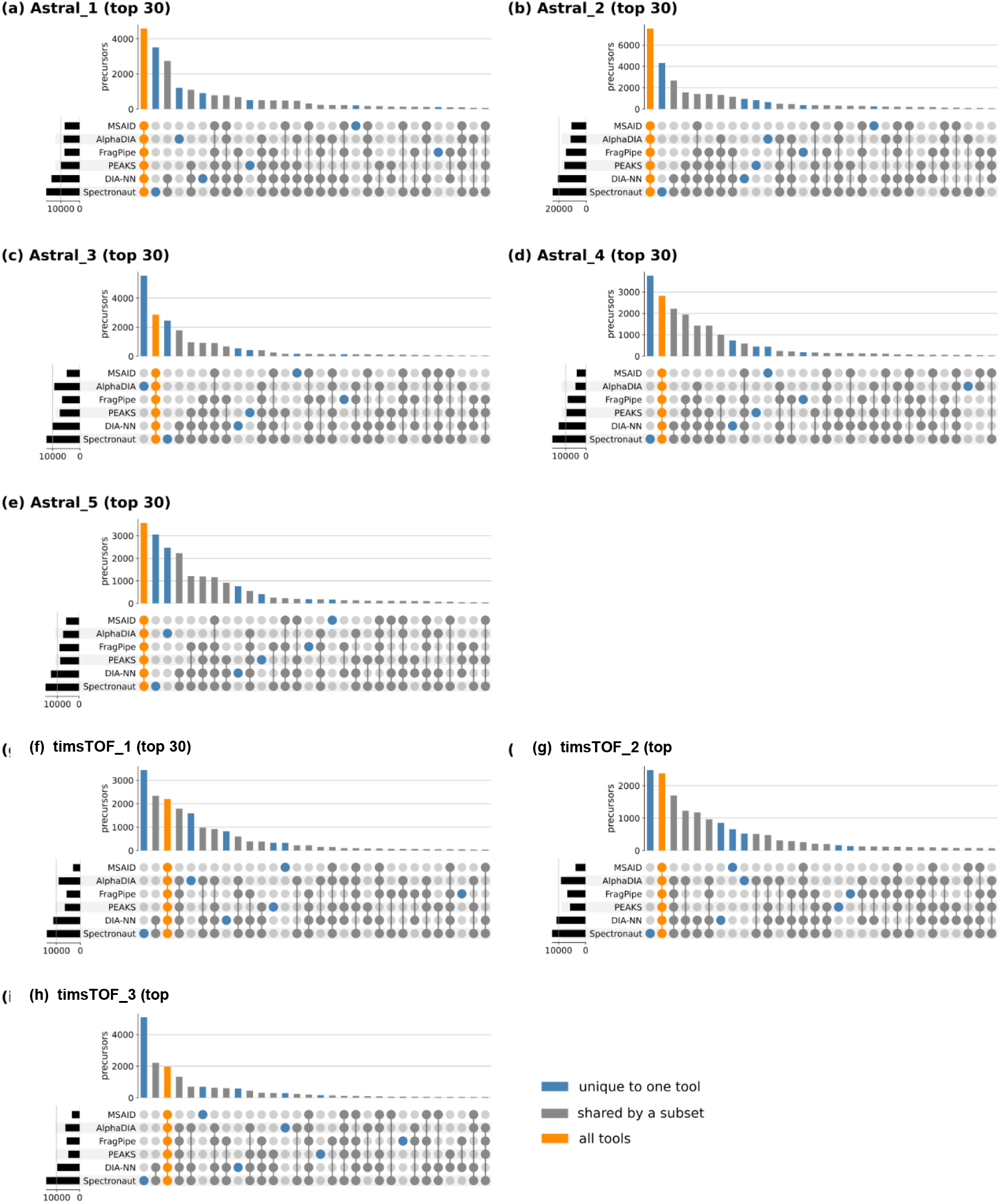
Peptide sequence overlap across all six search tools for **(a)** Astral_1, **(b)** Astral_3, **(c)** Astral_4, **(d)** Astral_5, **(e)** Astral_6, **(f)** timsTOF_1 and **(g)** timsTOF_3 instruments; blue bars indicate tool-unique identifications, orange indicates sequences identified by all six tools. Top 30 intersections are shown.

**Extended Figure 4:**
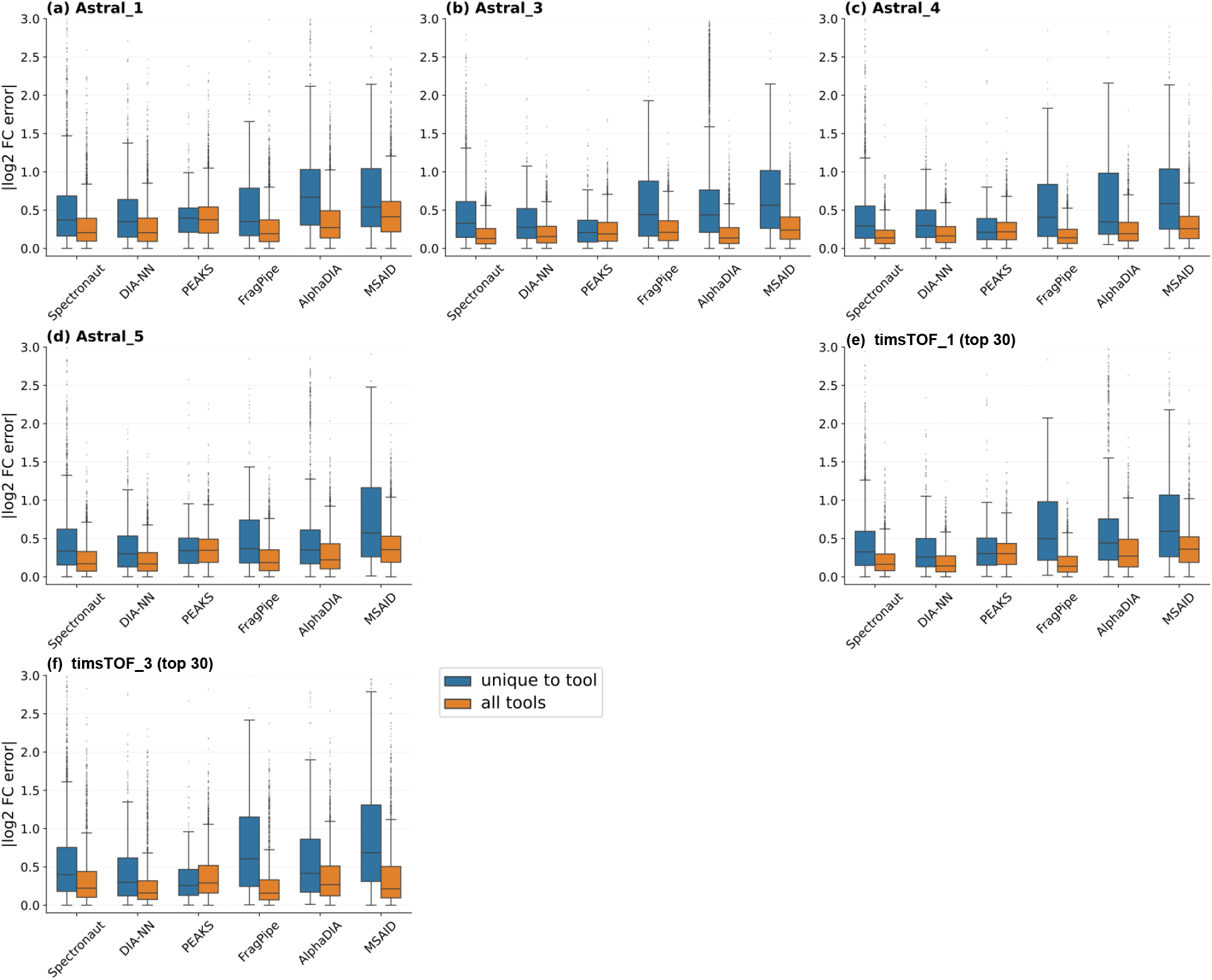
Absolute log_2_ fold-change error for peptides identified exclusively by a single tool (blue) versus peptides shared across all tools (orange) for **(a)** Astral_1, **(b)** Astral_3, **(c)** Astral_4, **(d)** Astral_5, **(e)** Astral_6, **(f)** timsTOF_1 and **(g)** timsTOF_3 instruments. Same as Fig. 2f-g for all other instruments.

**Extended Figure 5:**
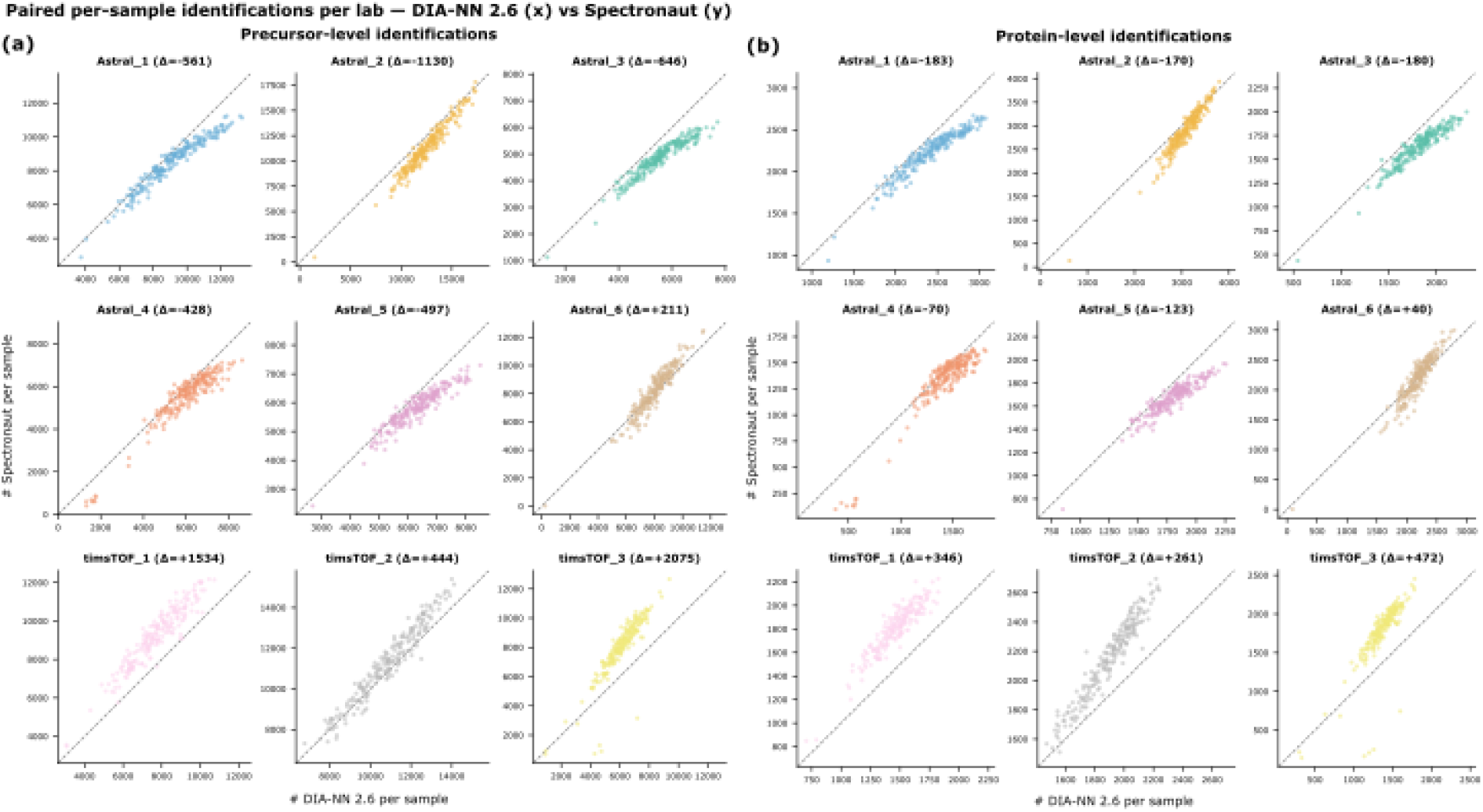
Each point represents one sample analyzed by both tools, restricted to the single-cell conditions (K562, HEK293T). The x-axis gives number of identifications reported by DIA-NN 2.6, y-axis Spectronaut 20. The dashed line marks equal detection; points above it indicate Spectronaut identified more features compared to DIA-NN. Each panel title reports median per-sample difference in identifications (Spectronaut - DIA-NN).

**Extended Figure 6:**
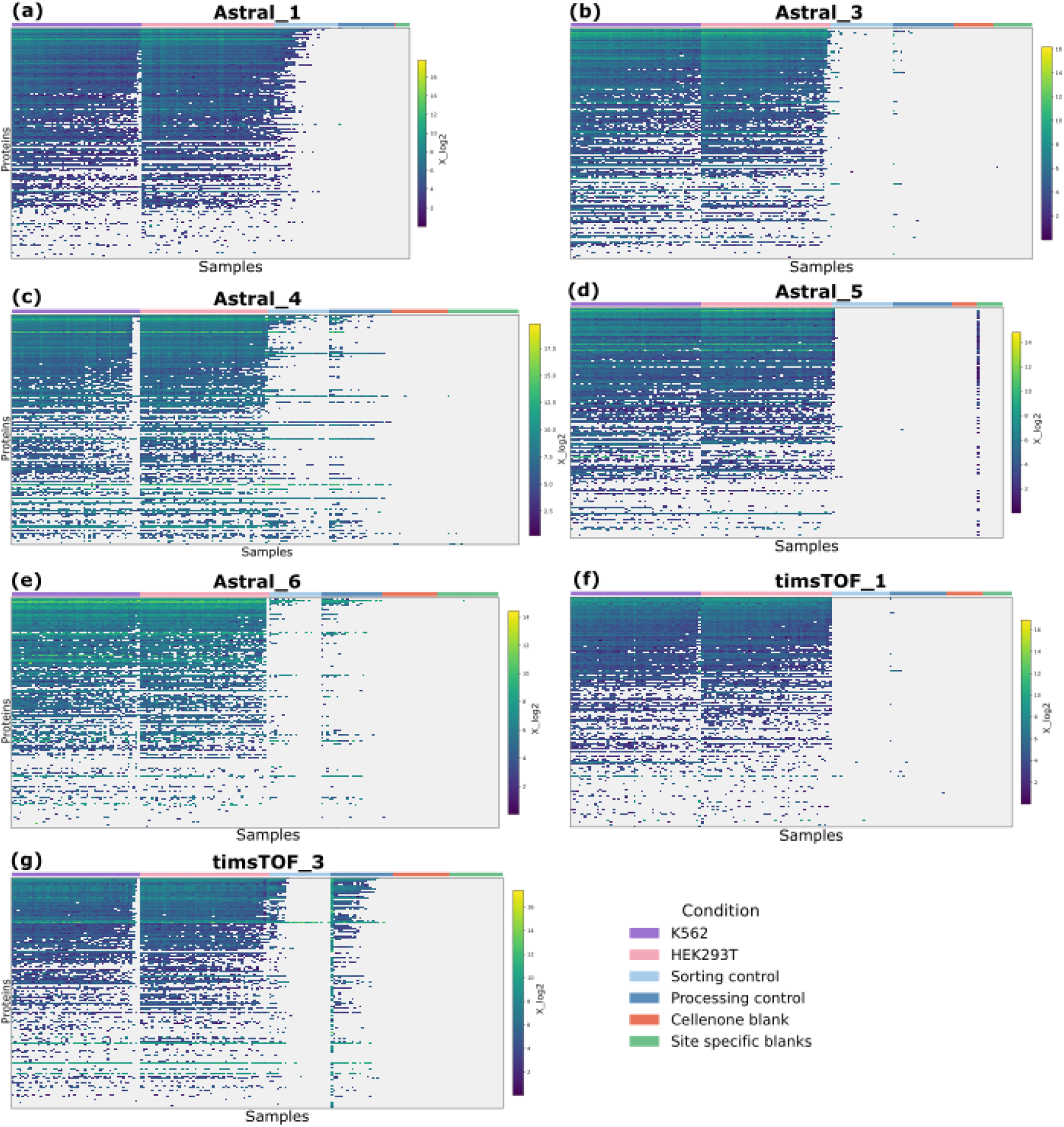
Overlap of protein identifications between single-cell samples (HEK293T, purple; K562, pink), processing controls (light blue), sorting controls (dark blue), and blank injections (orange and green) for **(a)** Astral_1, **(b)** Astral_3, **(c)** Astral_4, **(d)** Astral_5, **(e)** Astral_6, **(f)** timsTOF_1 and **(g)** timsTOF_3 instruments. Blue cells indicate identified proteins; grey cells indicate missing values. As Figure 3e-f for the other instruments.

**Extended Figure 7:**
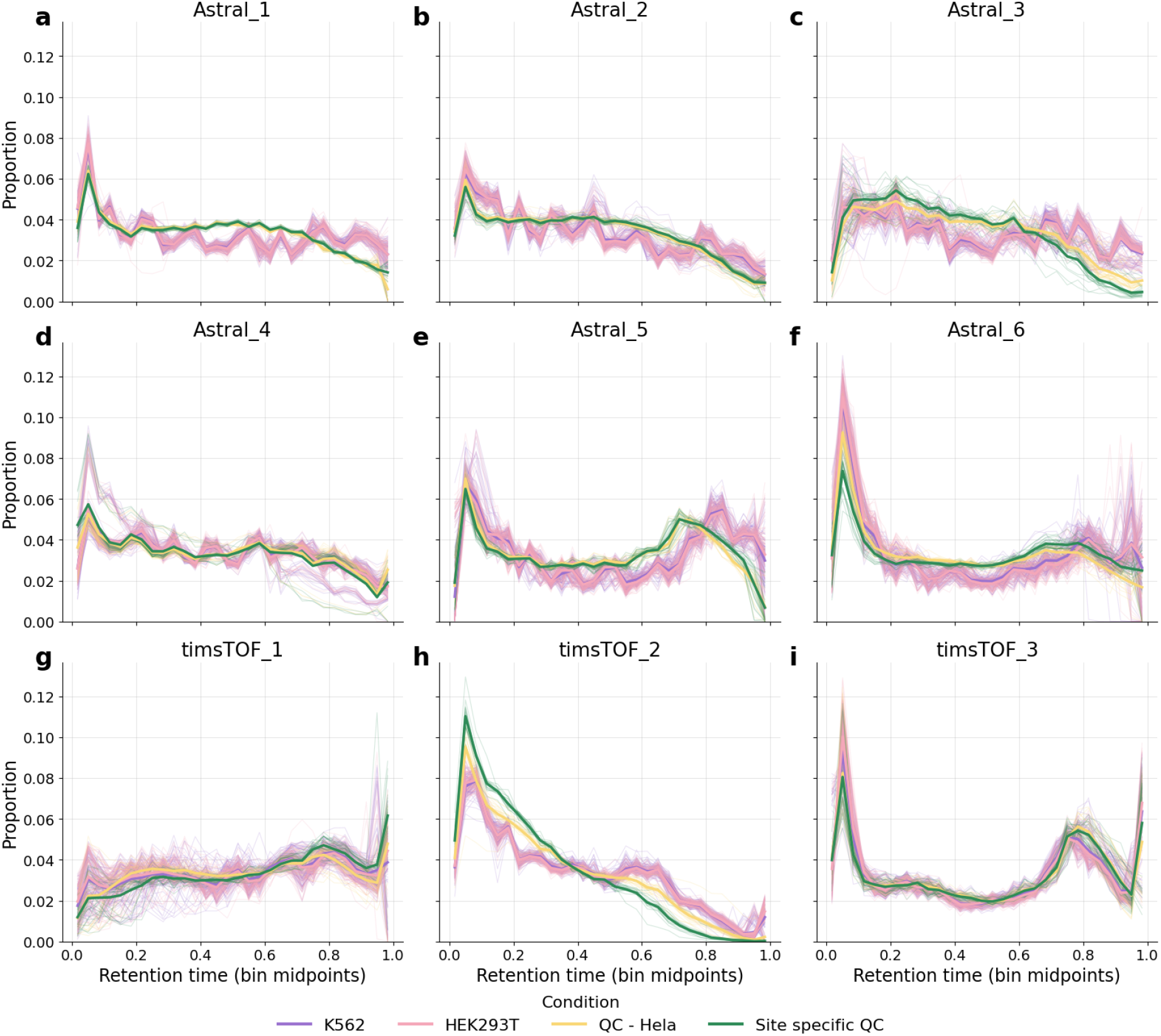
Proportion of precursors identified per RT bins, per sample one line is indicated where K562 is purple, HEK293T is pink, centralized QC is yellow, green is side-specific QC across all instruments across **(a)** Astral_1, **(b)** Astral_2, **(c)** Astral_3, **(d)** Astral_4, **(e)** Astral_5, **(f)** Astral_6, **(g)** timsTOF_1, **(h)** timsTOF_2 and **(g)** timsTOF_3 instruments.

**Extended Figure 8:**
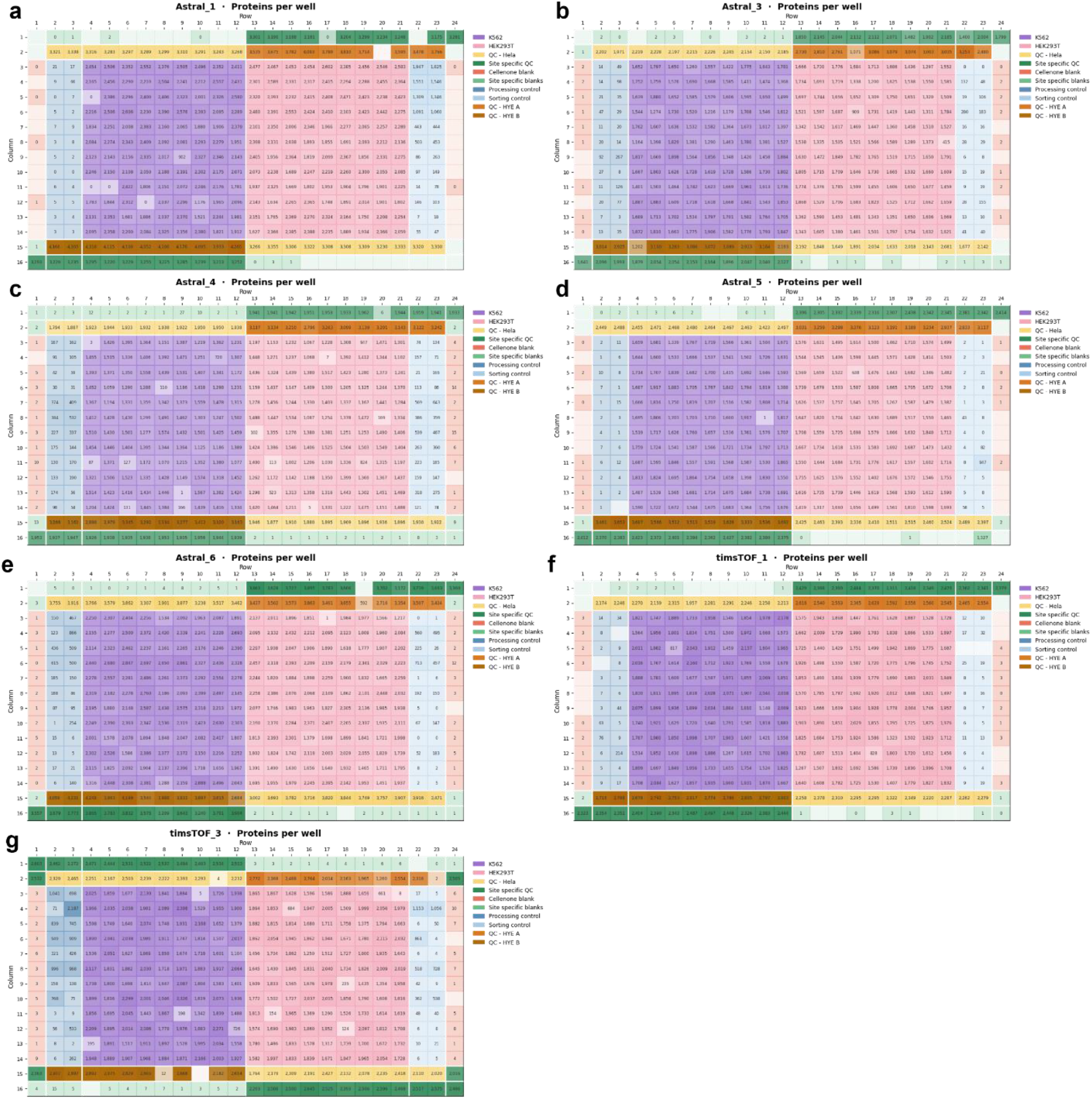
384-well plate overview showing precursor identifications per well for **(a)** Astral_1, **(b)** Astral_3, **(c)** Astral_4, **(d)** Astral_5, **(e)** Astral_6, **(f)** timsTOF_1 and **(g)** timsTOF_3 instruments; colors indicate sample group (K562 = purple, HEK293T = pink, centralized HeLa QC = yellow, site-specific HeLa QC = dark green, centralized blank = orange, site-specific blank = light green, quantitative QC = dark and light brown for Mix A or Mix B, processing control = dark blue, sorting control = light blue) and color intensity reflects relative identification counts within each group. As Figure 4b-c for all other instruments.

**Extended Figure 9:**
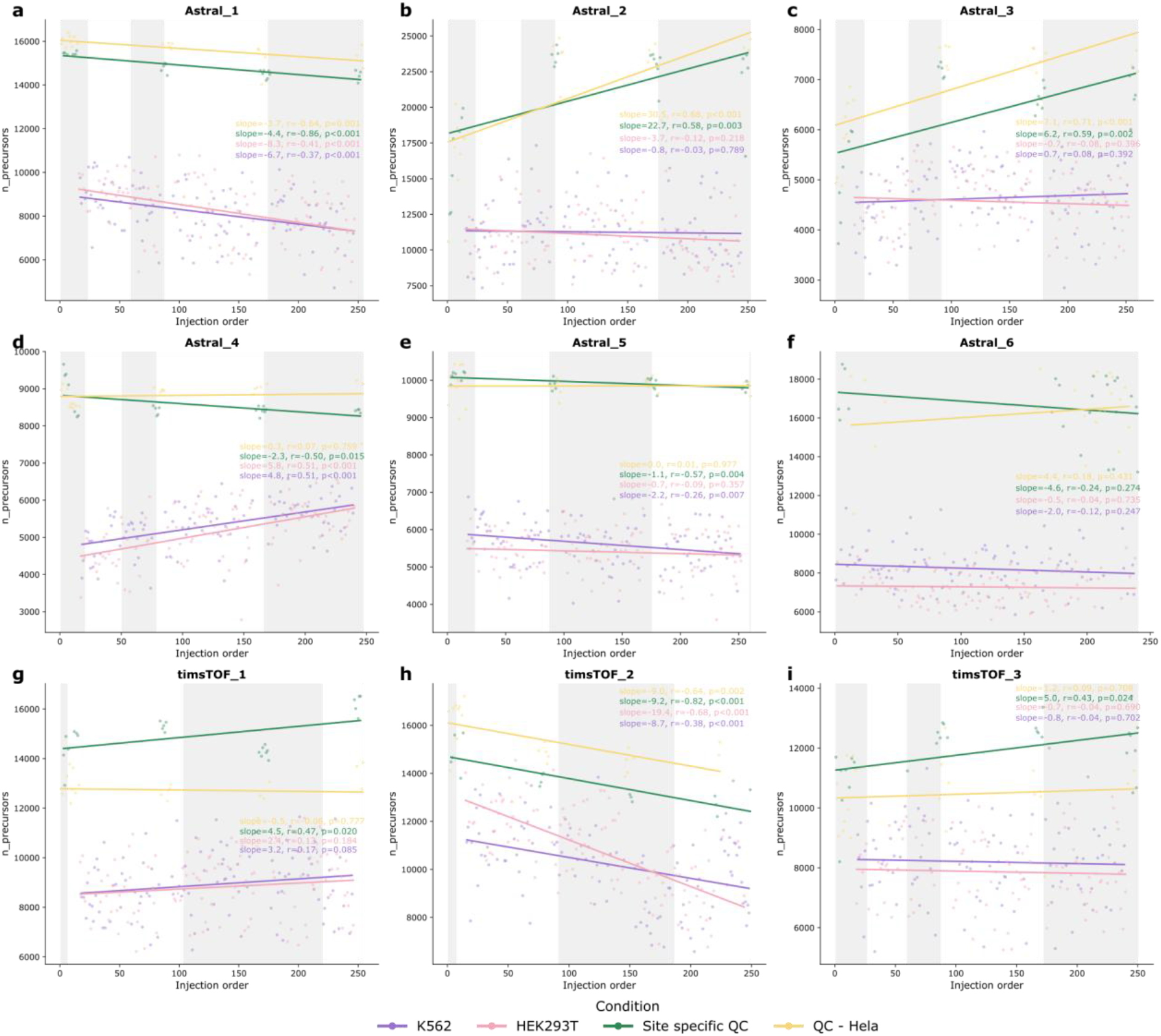
Boxplots showing the percentage of maximum precursor identifications per sample, binned by acquisition date, for K562 single cells, HEK293T single cells, centralized HeLa QC, and site-specific HeLa QC across all nine instruments: **(a)** Astral_1, **(b)** Astral_2, **(c)** Astral_3, **(d)** Astral_4, **(e)** Astral_5, **(f)** Astral_6, **(g)** timsTOF_1, **(h)** timsTOF_2, and **(i)** timsTOF_3. Shades of blue indicate successive acquisition dates and identification counts are normalized to the per-instrument maximum.

**Extended Figure 10:**
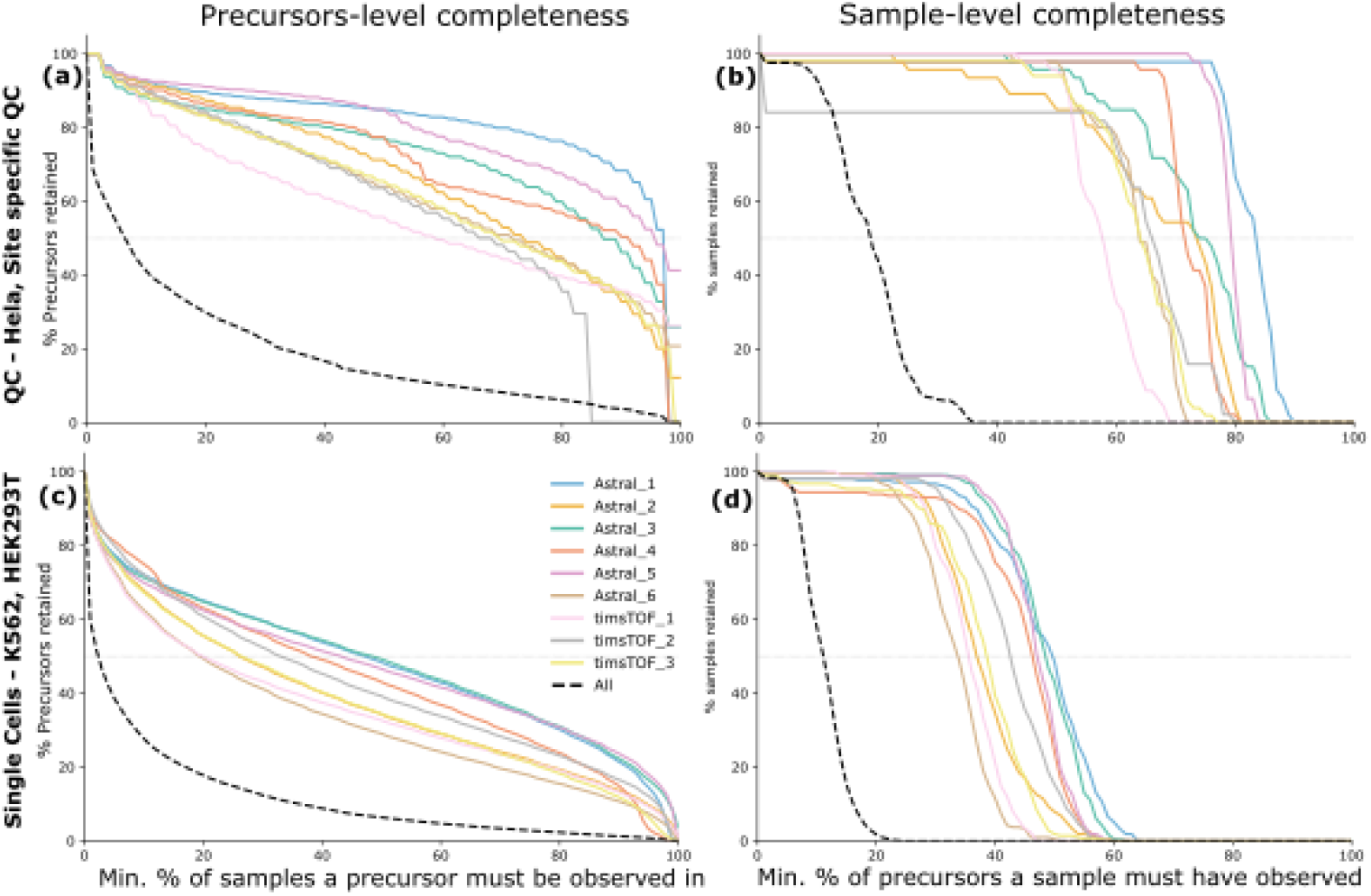
Feature-level data completeness showing the minimum percentage of samples in which a precursor must be observed versus the percentage of all precursors retained, for **(a)** single-cell samples and **(b)** HeLa QC samples. Sample-level data completeness showing the minimum percentage of precursors a sample must contain versus the percentage of samples retained, for **(c)** single-cell samples and **(d)** HeLa QC samples. Same as Figure 4e-h on precursor level.

## References

1. Budnik, B., Levy, E., Harmange, G. & Slavov, N. SCoPE-MS: mass spectrometry of single mammalian cells quantifies proteome heterogeneity during cell differentiation. Genome Biology 19, (2018).

2. Schoof, E. M. et al. Quantitative single-cell proteomics as a tool to characterize cellular hierarchies. Nat Commun 12, 3341 (2021).

3. Brunner, A.-D. et al. Ultra-high sensitivity mass spectrometry quantifies single-cell proteome changes upon perturbation. Molecular Systems Biology 18, e10798 (2022).

4. Zhu, Y. et al. Single-cell proteomics reveals changes in expression during hair-cell development. eLife 8, e50777 (2019).

5. Bubis, J. A. et al. Challenging the Astral mass analyzer - going beyond 5200 proteins per single-cell at unseen quantitative accuracy to study cellular heterogeneity. 2024.02.01.578358 Preprint at 10.1101/2024.02.01.578358 (2024).

6. Ctortecka, C. et al. Automated single-cell proteomics providing sufficient proteome depth to study complex biology beyond cell type classifications. Nat Commun 15, 5707 (2024).

7. Sabatier, P. et al. Global analysis of protein turnover dynamics in single cells. Cell 188, 2433–2450.e21 (2025).

8. Zhang, M. J., Ntranos, V. & Tse, D. Determining sequencing depth in a single-cell RNA-seq experiment. Nature Communications 11, 774 (2020).

9. Chen, W. et al. A multicenter study benchmarking single-cell RNA sequencing technologies using reference samples. Nature Biotechnology 1–12 (2020) doi:10.1038/s41587-020-00748-9.

10. Zhu, Y. et al. Nanodroplet processing platform for deep and quantitative proteome profiling of 10–100 mammalian cells. Nature Communications 9, 882 (2018).

11. Leduc, A., Huffman, R. G., Cantlon, J., Khan, S. & Slavov, N. Exploring functional protein covariation across single cells using nPOP. Genome Biology 23, 261 (2022).

12. Ye, Z. et al. One-Tip enables comprehensive proteome coverage in minimal cells and single zygotes. Nat Commun 15, 2474 (2024).

13. Lombard-Banek, C. et al. In Vivo Subcellular Mass Spectrometry Enables Proteo-Metabolomic Single-Cell Systems Biology in a Chordate Embryo Developing to a Normally Behaving Tadpole (X. laevis). Angewandte Chemie International Edition 60, 12852–12858 (2021).

14. Rodriguez, L. et al. Patch-Clamp Single-Cell Proteomics in Acute Brain Slices: A Framework for Recording, Retrieval, and Interpretation. 2025.09.15.675920 Preprint at 10.1101/2025.09.15.675920 (2025).

15. Cutler, R. et al. Mass spectrometry-based profiling of single-cell histone post-translational modifications to dissect chromatin heterogeneity. Nat Commun 16, 11100 (2025).

16. Matzinger, M., Müller, E., Dürnberger, G., Pichler, P. & Mechtler, K. Robust and Easy-to-Use One-Pot Workflow for Label-Free Single-Cell Proteomics. Anal. Chem. 95, 4435–4445 (2023).

17. Cong, Y. et al. Ultrasensitive single-cell proteomics workflow identifies >1000 protein groups per mammalian cell. Chemical Science https://doi.org/10.1039/D0SC03636F (2021) doi:10.1039/D0SC03636F.

18. Woo, J. et al. High-throughput and high-efficiency sample preparation for single-cell proteomics using a nested nanowell chip. Nat Commun 12, 6246 (2021).

19. Ctortecka, C. et al. An automated nanowell-array workflow for quantitative multiplexed single-cell proteomics sample preparation at high sensitivity. Molecular & Cellular Proteomics 0, (2023).

20. Liu, Z. et al. End-to-end high-throughput single-cell proteomics via SPRINT and dual-spray LC-MS. 2025.11.03.686420 Preprint at 10.1101/2025.11.03.686420 (2025).

21. Furtwängler, B. et al. Mapping early human blood cell differentiation using single-cell proteomics and transcriptomics. Science 0, eadr8785 (2025).

22. Su, P. et al. Proteoform profiling of endogenous single cells from rat hippocampus at scale. Nat Biotechnol 1–5 (2025) doi:10.1038/s41587-025-02669-x.

23. Wu, T. et al. Single-cell proteomic landscape of the developing human brain. Nat Biotechnol 1–14 (2026) doi:10.1038/s41587-025-02980-7.

24. Gatto, L. et al. Initial recommendations for performing, benchmarking and reporting single-cell proteomics experiments. Nat Methods 20, 375–386 (2023).

25. Yu, F. et al. Analysis of DIA proteomics data using MSFragger-DIA and FragPipe computational platform. Nat Commun 14, 4154 (2023).

26. Wallmann, G. et al. AlphaDIA enables DIA transfer learning for feature-free proteomics. Nat Biotechnol 1–10 (2025) doi:10.1038/s41587-025-02791-w.

27. Frejno, M. et al. Unifying the analysis of bottom-up proteomics data with CHIMERYS. Nat Methods 22, 1017–1027 (2025).

28. Demichev, V., Messner, C. B., Vernardis, S. I., Lilley, K. S. & Ralser, M. DIA-NN: neural networks and interference correction enable deep proteome coverage in high throughput. Nature Methods 17, 41–44 (2020).

