## SupplementalFigures for "Defining Quality Control Standards for Single-Cell Proteomics by Inter-Laboratory Benchmarking"

**Affiliations:**

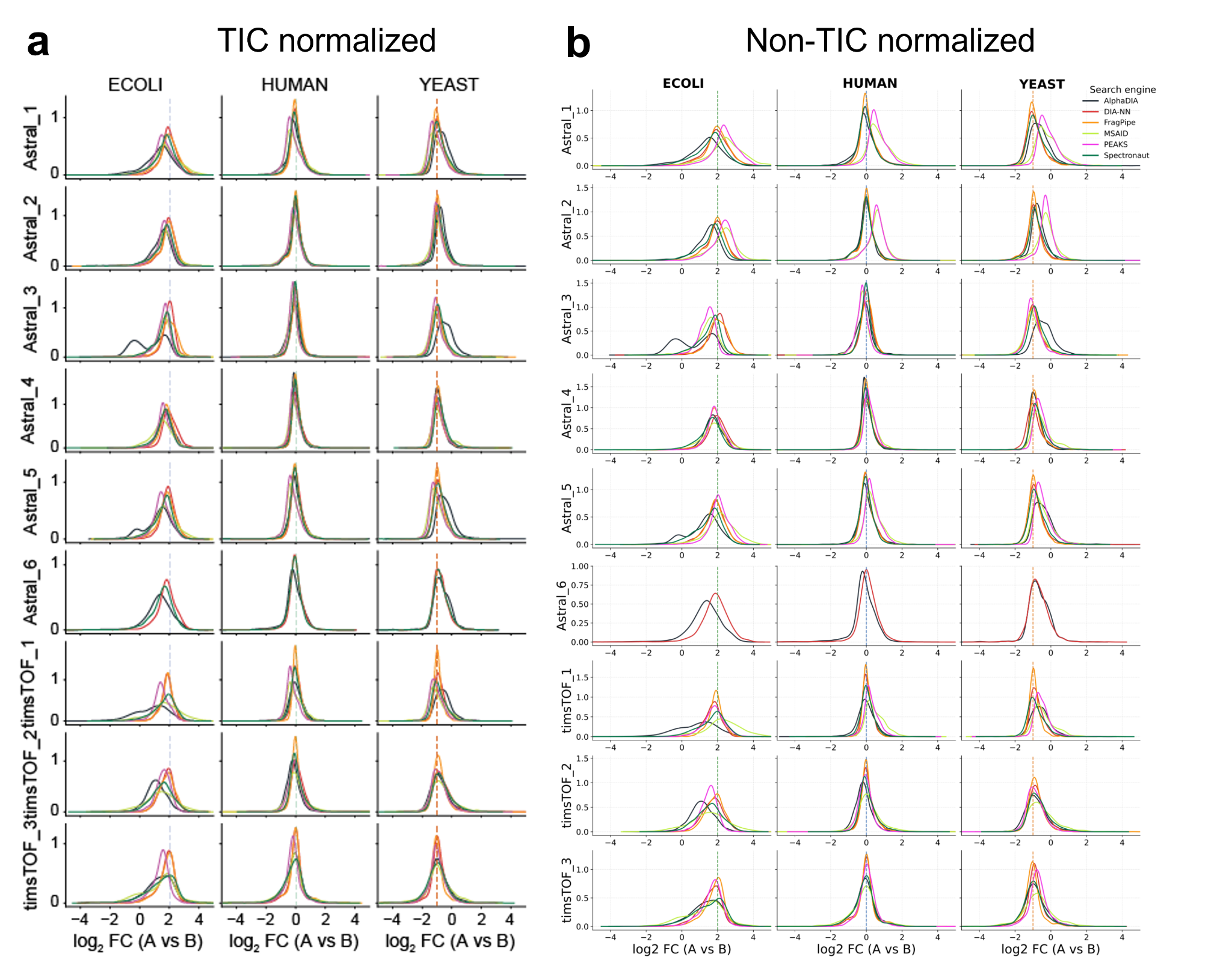

**Supplementary Figure 1: (a)** TIC normalized and **(b)** non-normalized log_2_ fold-change distributions for the three-proteome mixture separated by species is shown (left: E. coli; middle: human; right: yeast) with one row per instrument and colors indicate the different search tools.

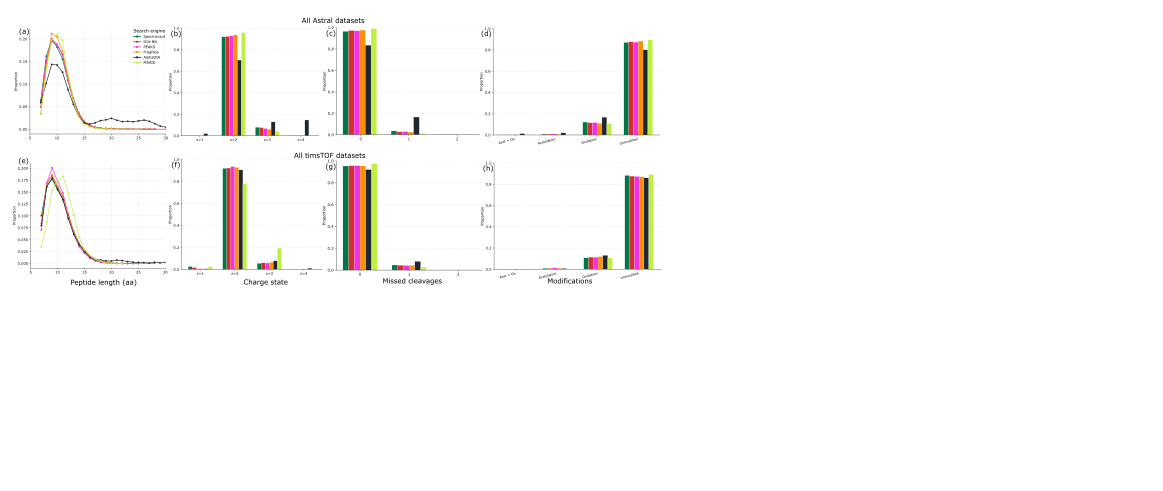

**Supplementary Figure 2:** Moieties of identified peptides across all search algorithms **(a)** peptide length, **(b)** peptide charge state, **(c)** missed cleavages, **(d)** post-translational modifications. Top row indicates Astral and bottom row timsTOF instruments.

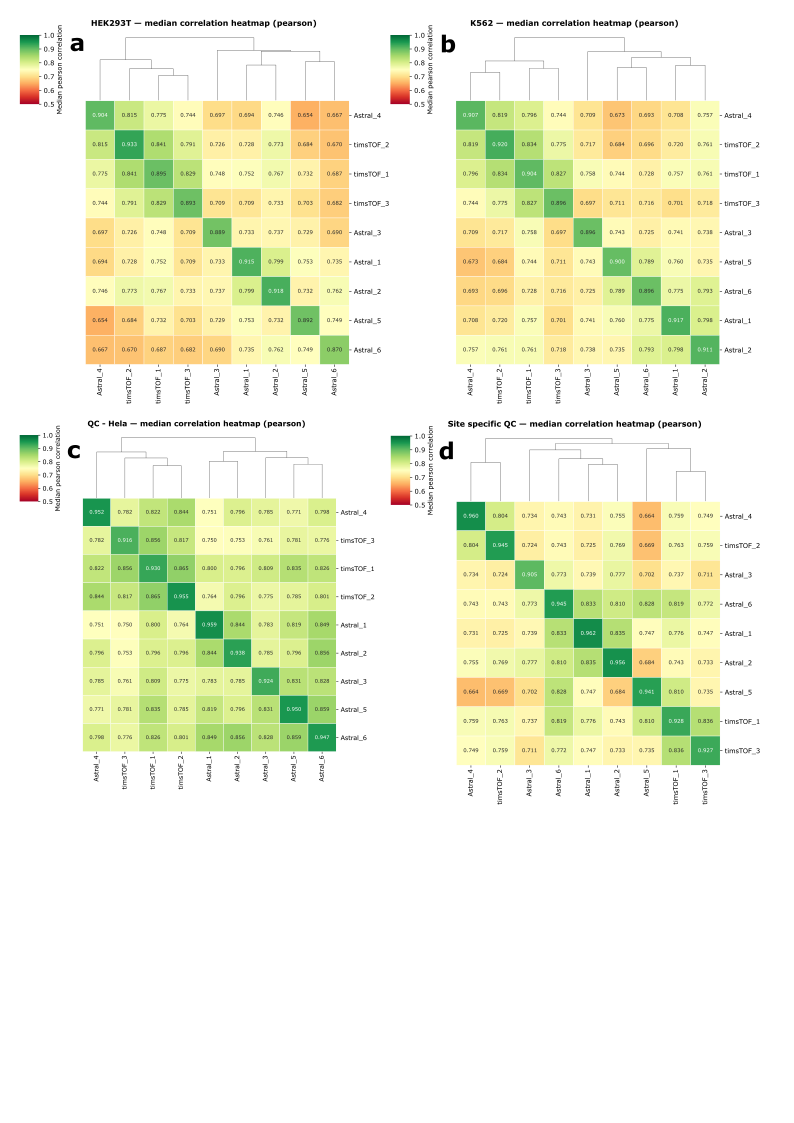

**Supplemental Figure 3:** Heatmaps showing the median pairwise Pearson correlation of precursor intensities between all instrument pairs for **(a)** HEK293T and **(b)** K562 single cells or **(c)** centralized and **(d)** site-specific HeLa QC. Color scale indicates Pearson correlation.

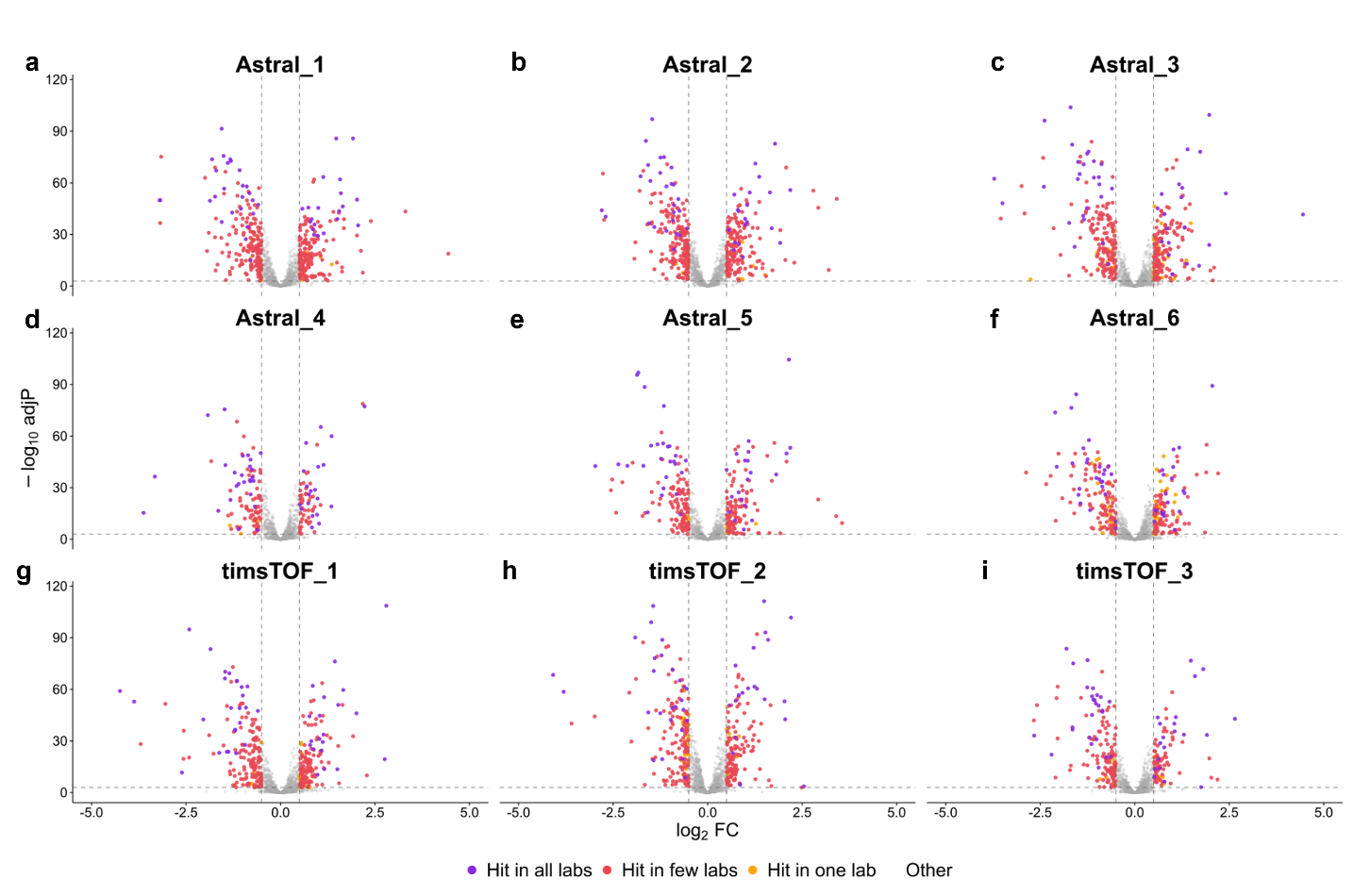

**Supplementary Figure 4:** Single-cell vulcano plots as shown in Fig. 6a labelled according to the frequency of identifications across multiple acquisition sites.
